# Discriminating Mild from Critical COVID-19 by Innate and Adaptive Immune Single-cell Profiling of Bronchoalveolar Lavages

**DOI:** 10.1101/2020.07.09.196519

**Authors:** Els Wauters, Pierre Van Mol, Abhishek D. Garg, Sander Jansen, Yannick Van Herck, Lore Vanderbeke, Ayse Bassez, Bram Boeckx, Bert Malengier-Devlies, Anna Timmerman, Thomas Van Brussel, Tina Van Buyten, Rogier Schepers, Elisabeth Heylen, Dieter Dauwe, Christophe Dooms, Jan Gunst, Greet Hermans, Philippe Meersseman, Dries Testelmans, Jonas Yserbyt, Patrick Matthys, Sabine Tejpar, CONTAGIOUS collaborators, Johan Neyts, Joost Wauters, Junbin Qian, Diether Lambrechts

## Abstract

How innate and adaptive lung immune responses to SARS-CoV-2 synchronize during COVID-19 pneumonitis and regulate disease severity is poorly established. To address this, we applied single-cell profiling to bronchoalveolar lavages from 44 patients with mild or critical COVID-19 *versus* non-COVID-19 pneumonia as control. Viral RNA-tracking delineated the infection phenotype to epithelial cells, but positioned mainly neutrophils at the forefront of viral clearance activity during COVID-19. In mild disease, neutrophils could execute their antiviral function in an immunologically ‘controlled’ fashion, regulated by fully-differentiated T-helper-17 (T_H17_)-cells, as well as T-helper-1 (T_H1_)-cells, CD8^+^ resident-memory (T_RM_) and partially-exhausted (T_EX_) T-cells with good effector functions. This was paralleled by ‘orderly’ phagocytic disposal of dead/stressed cells by fully-differentiated macrophages, otherwise characterized by anti-inflammatory and antigen-presenting characteristics, hence facilitating lung tissue repair. In critical disease, CD4^+^ T_H1_- and CD8^+^ T_EX_-cells were characterized by inflammation-associated stress and metabolic exhaustion, while CD4^+^ T_H17_- and CD8^+^ T_RM_-cells failed to differentiate. Consequently, T-cell effector function was largely impaired thereby possibly facilitating excessive neutrophil-based inflammation. This was accompanied by impaired monocyte-to-macrophage differentiation, with monocytes exhibiting an ATP-purinergic signalling-inflammasome footprint, thereby enabling COVID-19 associated fibrosis and worsening disease severity. Our work represents a major resource for understanding the lung-localised immunity and inflammation landscape during COVID-19.

## INTRODUCTION

SARS-CoV-2 has rapidly swept across the globe affecting >7 million people, with >400.000 fatal cases^1^. It is now well appreciated that while most COVID-19 patients (80%) remain asymptomatic or experience only mild symptoms, 20% present with overt pneumonia; about a quarter of these progress to a life-threatening state of Acute Respiratory Distress Syndrome (ARDS) and severe or atypical systemic inflammation^2^. Fever, increased acute phase reactants and coagulopathy with decreased lymphocyte counts, pronounced myeloid inflammation and increased neutrophil-to-lymphocyte ratio are predominant immunological hallmarks of severe COVID-19^3,4^.

Wen *et al.* were the first to provide an immune atlas of circulating mononuclear cells from 10 COVID-19 patients based on single-cell RNA-sequencing (scRNA-seq). Lymphocyte counts were globally decreased, while inflammatory myeloid cells, predominantly IL1β-secreting classical monocytes, were more abundant, suggesting COVID-19 immunopathology to be a myeloid-driven process^5^. Conversely, the contribution of circulating classical monocytes to systemic inflammation was put into question by Wilk *et al*. Based on scRNA-seq, they observed sparse expression of inflammatory cytokines by peripheral monocytes in 7 COVID-19 patients *versus* 6 six healthy controls. On the other hand, antigen presentation and the number of cytotoxic NK- and T-cells were reduced, while plasmablasts and neutrophils were increased, especially in in COVID-19 patients experiencing ARDS^6^.

However, profiling the peripheral immune landscape in COVID-19 may not be as comprehensive since immune characteristics in the periphery are different from those within the lungs, both in terms of amplitude and qualitative characteristics, as well as duration of the immune response. Thus, a better understanding of the immune interactions in COVID-19 lungs is needed. In their seminal paper, Liao et al. applied single-cell T-cell receptor-sequencing (scTCR-seq) and scRNA-seq on bronchoalveolar lavage (BAL) fluid from 3 mild and 6 critical COVID-19 patients, as well as 3 healthy controls. They observed an abundance of highly inflammatory monocytes and neutrophils and T-cell depletion in critical *versus* mild COVID-19. The latter showed a more potent adaptive immune response to SARS-CoV-2, evidenced by presence of CD8^+^ T-cells with tissue-resident features displaying clonal expansion and increased effector function^7^. Subsequently, Bost *et al.* were able to sort infected cells from bystander cells and investigate virus-induced transcriptional changes. This Viral-Track pipeline showed ciliated and progenitor epithelial cells to be the main targets of SARS-CoV-2, yet a strong enrichment of viral RNA was observed in SPP1+ macrophages^8^. It is unclear however whether this represents direct viral infection of myeloid cells, or phagocytosis of viral particles (or virus-infected cells). Moreover, due to the small sample size, in-depth characterization of all the cellular phenotypes detected by scRNA-seq in mild *versus* critical COVID-19 remained largely unexplored.

Here, we provide a comprehensive deep-immune atlas of COVID-19 pneumonitis, analyzing BAL from 31 COVID-19 patients with mild or critical disease, while inclusion of 13 patients with non-COVID-19 pneumonitis allowed us to reliably distinguish non-specific lung-localised inflammatory signaling from COVID-19 specific lung-associated immune changes.

## RESULTS

### scRNA-seq and cell typing of BAL samples

We performed scRNA-seq on BAL from 22 hospitalized patients with a positive qRT-PCR for SARS-CoV-2 on a nasopharyngeal swab or a lower respiratory tract sample. We also collected BAL from 13 patients with clinical suspicion of COVID-19 pneumonia, yet negative PCR on lower respiratory tract sampling for SARS-CoV-2. These samples are referred to as non-COVID-19. We further stratified COVID-19 patients by disease severity at the time of sampling, by discerning two groups; a ‘mild’ (n=2) and a ‘critical’ (n=20) disease group, the latter requiring mechanical ventilation or extracorporeal membrane oxygenation. Demographic and clinical data of the prospectively recruited patient cohort are summarised in Supplementary information, **Table S1**.

BAL samples were immediately processed for scRNA-seq. After quality filtering (Methods), we obtained ~186 millions of unique transcripts from 65,166 cells with >150 genes detected. Of these, ~51% of cells were from COVID-19. Subsequent analysis involving dimensionality reduction and clustering identified several clusters (**Fig. 1a**), which through marker genes (Supplementary information, **Fig. S1a, b**) could be assigned to lung epithelial cells (including ciliated, inflammatory, hillock, secretory and AT2 lung epithelial cells), myeloid cells (monocytes/macrophages, neutrophils, mast cells, plasmacytoid dendritic cells/pDCs and conventional dendritic cells/cDCs), lymphoid cells (CD4^+^ and CD8^+^ T-cells, natural killer cells (NK), B-cells and plasma cells). We describe each cell type in more detail, highlighting the number of cells, read counts and transcripts detected in Supplementary information, **Table S2**. There was no cluster bias between disease status (COVID-19 *versus* non-COVID-19), disease severity (mild *versus* critical) or individual patients (**Fig. 1b**).

**Fig. 1.**
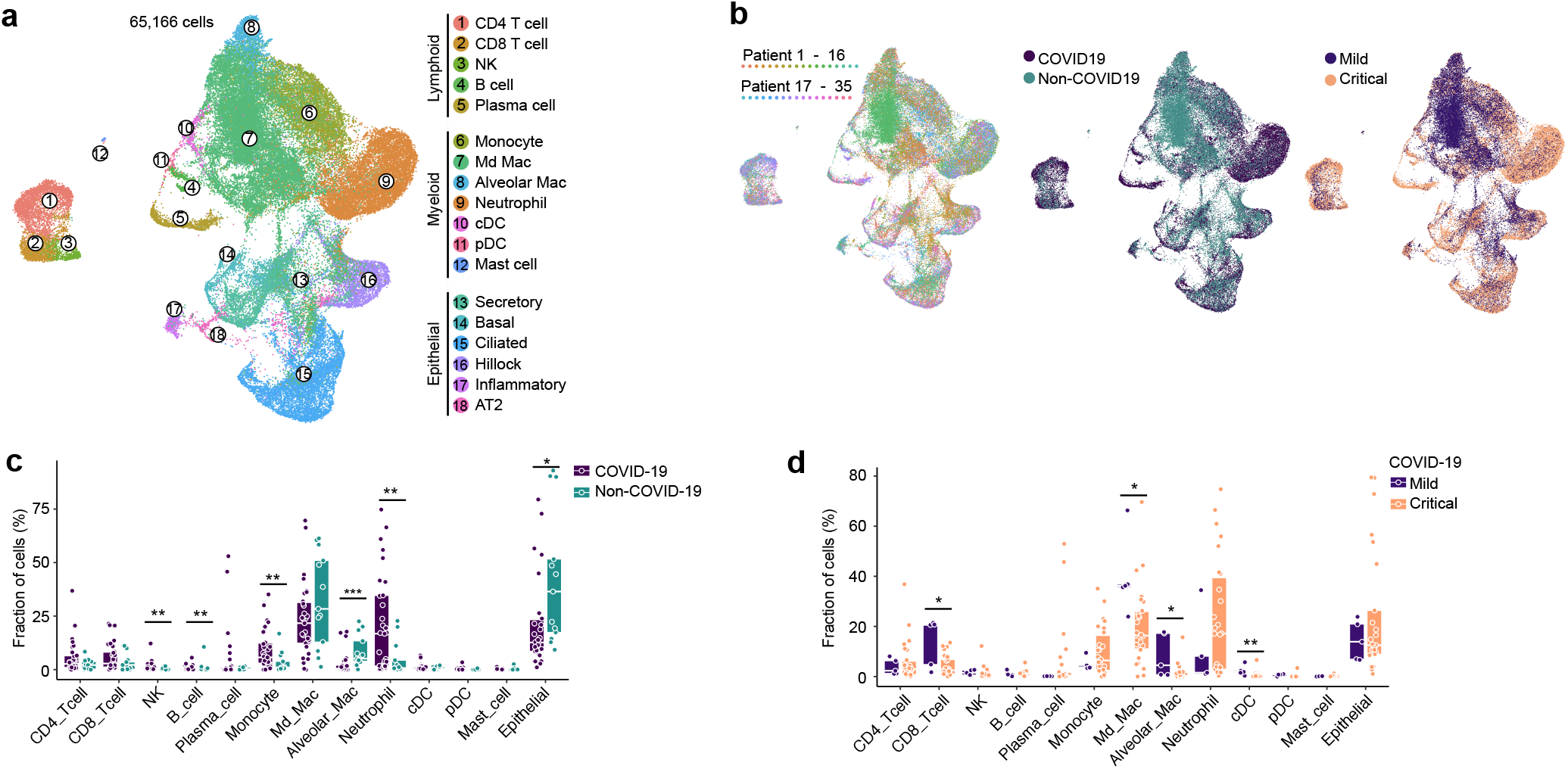
Annotation of cell types by scRNA-seq in COVID-19 and non-COVID-19 BAL. **a** UMAP representation of 65,166 cells (obtained from BAL from n=13 non-COVID-19, n=2 mild and n=22 critical COVID-19 patients) by scRNA-seq color-coded for the indicated cell type. pDC: plasmacytoid dendritic cell, cDC: conventional dendritic cell, NK: natural-killer cell, Md_Mac: monocyte-derived macrophage. Alveolar_Mac: alveolar macrophage. AT2: alveolar type II epithelial cell. **b** UMAP panels stratified per individual patient, COVID-19 *versus* non-COVID-19 and mild *versus* critical COVID-19. **c** Relative contribution of each cell type (in %) in COVID-19 *versus* non-COVID-19. **d** Relative contribution of each cell type (in %) in mild *versus* critical COVID-19. P values were assessed by Mann-Whitney test. * P<0.05, ** P<0.01, ***P<0.001

To increase our resolution, we processed scRNA-seq data on COVID-19 BAL by Liao et al., consisting of 3 patients with ‘mild’ and 6 patients with ‘critical’ COVID-19 (n=51,631 cells) (Supplementary information, **Fig. S1c**)^7^. We also retrieved 7 normal lung samples (n=64,876 cells) profiled by Lambrechts et al. and 8 normal lung samples (n= 27,266 cells) by Reyfman et al. to further enhance our resolution, specifically for T-cells and DCs^9,10^. Datasets were integrated by clustering cells from each dataset separately and assigning cell type identities to each cell. We then pooled cells from each dataset based on cell type identities and performed canonical correlation analysis (CCA), as described previously^11^, followed by graph-based clustering to generate a UMAP per cell type, displaying its phenotypic heterogeneity. Per cell type, we ruled out batch effects due to different datasets.

After integration, COVID-19 BAL scRNA-seq data were derived from 5 mild and 26 critical COVID-19 patients. Quantitatively, monocyte/macrophages and neutrophils were the most abundant cell types, amounting up to 65.7% (n=55,825) of COVID-19 cells (**Fig. 1c**). When evaluating the relative enrichment or depletion of these cell types, we found that monocytes and neutrophils were indeed more frequent, while macrophages and epithelial cells were less abundant in COVID-19 *versus* non-COVID-19. B-cells and NK-cells were slightly enriched in COVID-19. Comparing mild *versus* critical COVID-19 revealed an increase in CD8^+^ T-cells, macrophages and cDCs in the former (**Fig. 1d**).

Below, we describe the heterogeneity underlying each cell type in more detail.

### Phenotypic heterogeneity of CD8^+^ T-cells in COVID-19 BAL

Altogether, we retrieved 23,468 T- and NK-cells, which were subclustered into 14 phenotypes (**Fig. 2a, b**; Supplementary information, **Fig. S2**). Briefly, we identified 7 CD8^+^ T-cell clusters, 5 CD4^+^ T-cell clusters and 2 NK-cell clusters. While naïve CD8^+^ T-cells (T_N_) expressed naïve T-cell markers (*CCR7*, *LEF1* and *TCF7*), effector-memory (T_EM_) and partially-exhausted (T_EX_) T-cells were characterized by increased expression of effector markers (*IFNG*, *PRF1*, *NKG7* and *GZMB*). Herein, expression of (inflammation-driven) exhaustion-defining immune-checkpoints (*LAG3*, *HAVCR2* and *PDCD1*) distinguished T_EX_-cells. Additionally, we identified CD8^+^ resident-memory T-cells (T_RM_) based on *ZNF683* and *ITGAE*, as well as CD8^+^ recently-activated effector-memory T-cells (T_EMRA_). Finally, we also identified mucosal-associated invariant T-cells (T_MAIT_) and gamma-delta (T_γδ_) T-cells.

**Fig. 2.**
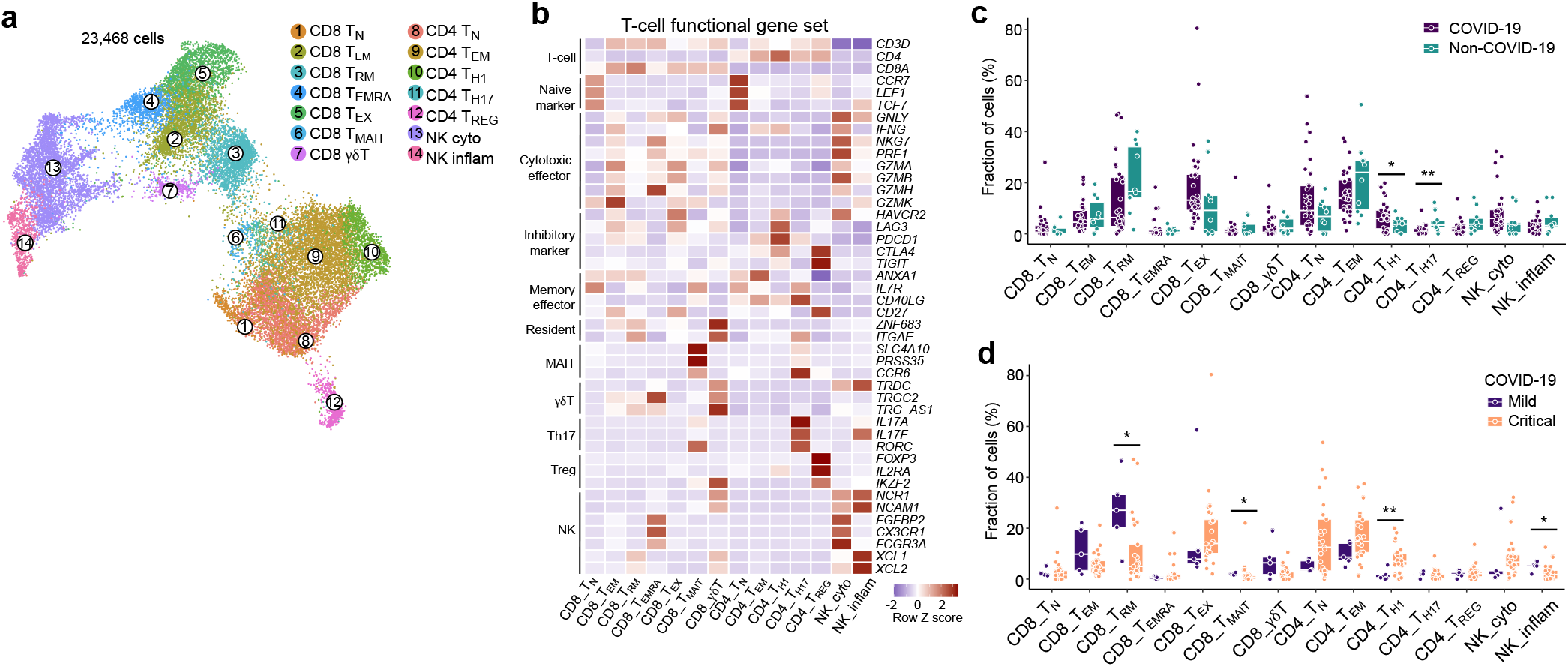
14 T-cell phenotypes in mild and critical COVID-19 BAL. **a** Subclustering of 23,468 T-/NK-cells into 14 T-/NK-cell phenotypes, as indicated by the color-coded legend. NK cyto: cytotoxic NK cell; NK inflam: inflammatory NK cell. **b** Heatmap showing T-/NK-cell phenotypes with corresponding marker genes and functional gene sets. **c** Relative contribution of each T-/NK-cell phenotype (in %) in COVID-19 *versus* non-COVID-19. **d** Relative contribution of each T-/NK-cell phenotype (in %) in mild *versus* critical COVID-19. P values were assessed by Mann-Whitney test. * P<0.05, ** P<0.01, ***P<0.001

Next, we assessed prevalence of each T-cell phenotype in COVID-19 *versus* non-COVID-19 disease, but failed to observe differences in the CD8^+^ phenotypes (**Fig. 2c**). When comparing mild to critical COVID-19 (**Fig. 2d**), we found T_MAIT_-cells to be increased in the former. Interestingly, T_MAIT_-cells can actively co-opt for specific innate immune characteristics (e.g. proficient pattern-recognition receptor-based signalling, and/or broad non-MHC antigenic surveillance), thereby allowing them to rapidly respond to pathogenic agents possessing pathogen-associated molecular patterns (PAMPs)^12^.

The largest increase in mild *versus* critical COVID-19, however, was seen for CD8^+^ T_RM_-cells. To understand this difference, we used Slingshot to infer pseudotime trajectories (excluding CD8^+^ T_MAIT_- and T_γδ_-cells). We observed 3 distinct CD8^+^ T-cell trajectories (**Fig. 3a**): CD8^+^ T_N_-cells connected with T_EM_-cells, which subsequently branched into 3 different (well-connected) lineages i.e., T_RM_-cells, T_EX_-cells and T_EMRA_-cells. Profiling of marker genes along these trajectories confirmed their functional annotations (**Fig. 3b**). Notably, effector function peaked halfway in each lineage and then stabilized or decreased, depending on the lineage (**Fig. 3c**). Next, density plots reflecting the relative number of T-cells in each phenotypic state were created along these trajectories (**Fig. 3d**), and stratified for normal tissue, non-COVID-19, and mild or critical COVID-19. Non-COVID-19 T-cells were enriched towards the end of the T_RM_-lineage, while in COVID-19 they were more frequent in the T_EX_-lineage (**Fig. 3e**). In contrast, T_EMRA_-cells were more differentiated in normal lung. When comparing mild to critical disease, the T_RM_-lineage was more differentiated in mild COVID-19, while along the T_EX_-lineage differentiation was most prominent in critical COVID-19 (**Fig. 3f**). There were no differences in the T_EMRA_-lineage.

**Fig. 3.**
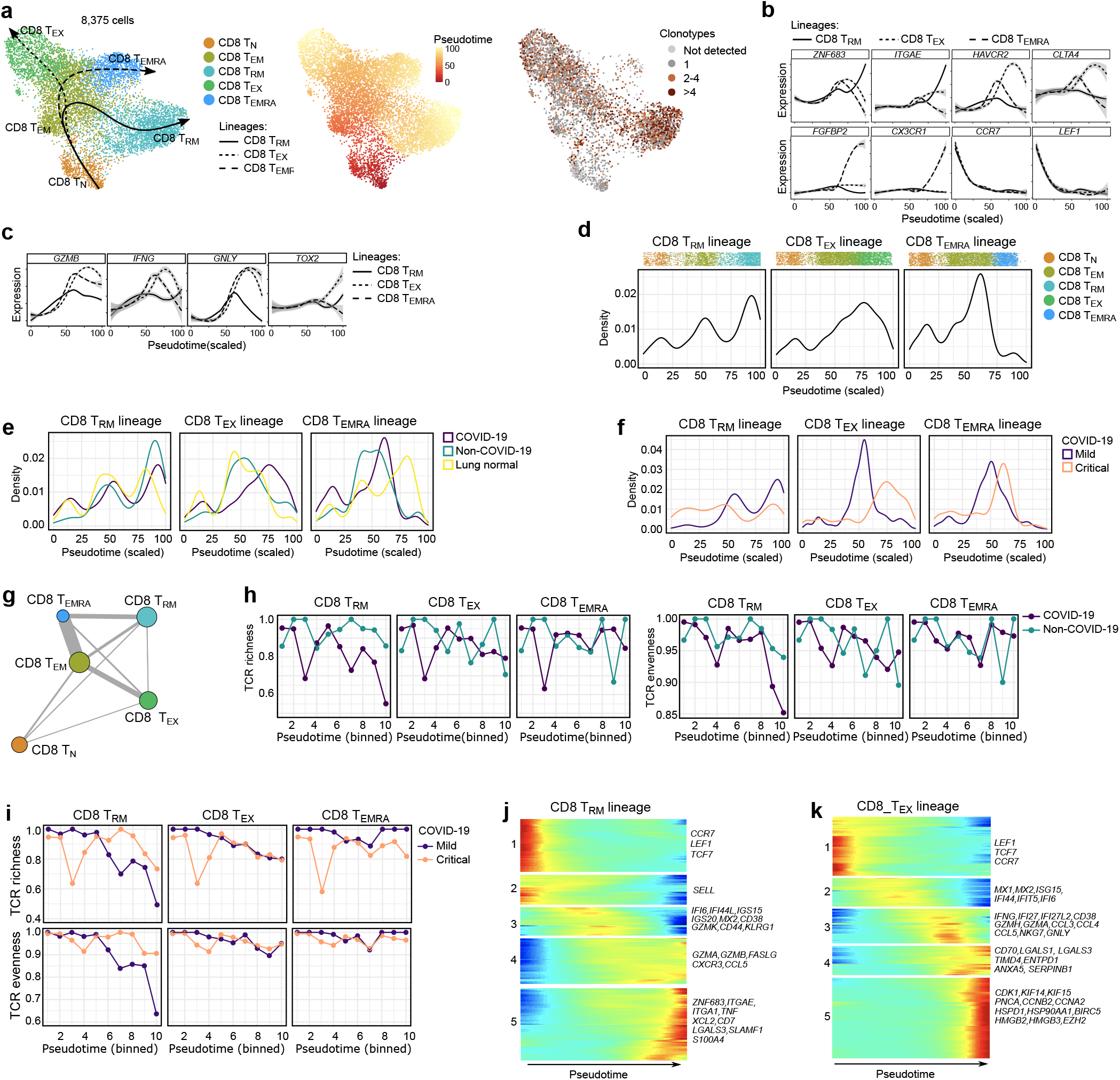
CD8^+^ T-cell phenotypes in mild and critical COVID-19 BAL. **a** Pseudotime trajectories for CD8^+^ T-cells based on Slingshot, showing 3 lineages (T_RM_-lineage, T_EX_-lineage and T_EMRA_-lineage), color-coded for the CD8^+^ T-cell phenotypes (left panel), the pseudotime (middle panel) and the number of clonotypes (right panel). **b** Profiling of marker genes along these trajectories to confirm their functional annotation: *ZNF683* and *ITGAE* for the T_RM_-lineage, *HAVCR2* and *CTLA4* for the T_EX_-lineage, *FGFBP2* and *CX3CR1* for the T_EMRA_-lineage. **c** Genes involved in T-cell effector function and cytotoxicity (*GZMB*, *IFNG*, *GNLY*) and related transcription factor (*TOX2*) modelled along the CD8^+^ T-cell lineages. **d** Density plots reflecting the number of T-cells along the 3 CD8^+^ T-cell lineages. **e** Density plots reflecting the number of T-cells along the 3 CD8^+^ T-cell lineages stratified for non-COVID-19, COVID-19 and normal lung. **f** Density plots reflecting the number of T-cells along the 3 CD8^+^ T-cell lineages stratified for mild *versus* critical COVID-19. **g** Analysis of clonotype sharing (thickness indicates proportion of sharing) between the CD8^+^ T-cells. **h-i** TCR richness and TCR evenness along the 3 T-cell lineages for non-COVID-19 *versus* COVID-19 (**h**), and mild *versus* critical COVID-19 (**i**). **j-k** Gene expression dynamics along the CD8^+^ T_RM_-(**j**) and T_EX_-lineage (**k**). Genes cluster into 5 gene sets, each of them characterized by specific expression profiles, as depicted by a selection of marker gene characteristic for each set. Differences in trajectories were assessed by Mann-Whitney test. For CD8^+^ T_RM_: COVID-19 *versus* non-COVID-19 (P =1.0E-6), mild *versus* critical COVID-19 (P=5.9E-102). For CD8^+^ T_EX_: COVID-19 *versus* non-COVID-19 and normal lung (P=2.3E-67), mild *versus* critical (P=1.1E-39). For CD8^+^ T_EMRA_: normal lung *versus* COVID-19 and non-COVID-19 (P=3.8E-39).

We also processed T-cells by scTCR-seq, obtaining 3,966 T-cells with a TCR sequence that were also annotated by scRNA-seq (excluding NK-, T-_MAIT_ and T_γδ_-cells). Based on TCR sharing, we could reinforce the 3 trajectories identified by Slingshot (**Fig. 3g**). Overall, CD8^+^ T_RM_-cells contained the highest number of T-cell clonotypes. Plotting TCR richness and evenness along the trajectories, revealed that both parameters were reduced along the T_RM_-lineage, specifically in COVID-19 (**Fig. 3h**), likely indicating antigen-driven clonal expansion. Notably, this expansion was more prominent in mild COVID-19 (**Fig. 3i**). In contrast, T_EX_-cells were characterized by only a modest decrease in TCR richness/evenness along their lineage. In the T_EMRA_ lineage, richness did not decrease along the pseudotime.

Overall, this suggests that mild COVID-19 is characterized by fully-differentiated T_RM_-cells undergoing active (presumably antigen-driven) TCR expansion and selection, while T_EX_-cells are entangled halfway their trajectory. In critical COVID-19, T_RM_-cells fail to differentiate or expand, while T_EX_-cells become more exhausted albeit without undergoing clonal expansion.

### Gene expression modelling along the CD8^+^ T_RM_- and T_EX_-lineage

We then modelled gene expression along the T_RM_- and T_EX_-lineage, and identified 5 gene sets with a specific expression pattern in each trajectory. In the T_RM_-lineage, set 1 and 2 consisted of naïve T-cell markers (set 1: *CCR7*, *LEF1, TCF7*; set 2: *SELL*), whose expression decreased along the trajectory (**Fig. 3j**; Supplementary information, **Table S3**). A third set was enriched for interferon (IFN)-induced (anti-viral) genes (*IFI6*, *IFI44L*, *ISG15*, *ISG20*, *MX2*), activation-associated genes (*CD38*) and genes mediating effector-memory functions (*GZMK, CD44, KLRG1*). Set 3 exhibited high expression halfway of the trajectory. Genes from the last 2 sets were expressed at the end of the trajectory and were mostly related to cytotoxicity and increased effector function (set 4: *GZMA, GZMB, FASLG, CXCR3, CCL5*), a balance of pro-inflammatory and auto-regulatory genes (set 5: *ITGA1*, *TNF*, *XCL2, CD7 versus LGALS3*, *SLAMF1*, *S100A4*) and genes marking resident memory formation (*ZNF683, ITGAE*)^13,14^. In mild COVID-19, T_RM_-cells mainly expressed set 3-5 genes, indicating increased (but balanced) effector function.

In the T_EX_-lineage, the first set contained naïve markers (*LEF1*, *CCR7*, *TCF7*), a second set IFN-induced (anti-viral) genes (*MX1*, *MX2, ISG15*, *IFI44*, *IFIT5*, *IFI6*), while a third set besides *IFNG* and IFN-induced genes (*IFI27*, *IFI27L2*) was also comprised of T-cell activation-related genes (*CD38*, *GZMH*, *GZMA*), chemokines (*CCL3*, *CCL4* and *CCL5*), cytotoxicity-(*NKG7*, *GNLY*, *GZMB*) and (inflammatory) exhaustion-related genes (*HAVCR2*) (**Fig. 3k**; Supplementary information, **Table S4**). Set 4 was characterized by expression of pro-inflammatory (*CD70*, *COTL*, *HMGB1*) and anti-inflammatory genes (*ENTPD1*, *ANXA5*, *SERPINB1*), suggesting these cells exhibit a chronic dysregulated hyperinflammatory phenotype. We also noticed expression of the TIM auto-regulatory protein family (*TIMD4*) and viral infection-induced auto-regulatory genes (*LGALS1*, *LGALS3*)^15,16^. In set 5, cell-cycle genes (*CDK1*, *KIFs*, *PCNA*, *CCNA/B2*), stress-associated genes (*HSPD1*, *HSP90AA1*, *BIRC5*) and chromatin re-modelling related genes (e.g., *HMGB2*, *HMGB3*, *EZH2*) were increased, suggesting that T-cells were largely adjusting to inflammation-driven stress (rather than mounting any discernible effector or auto-regulatory responses). Notably, mild COVID-19 showed a very prominent enrichment in cells characterized by set 3-associated genes, whereas critical COVID-19 was enriched for set 4-5 T_EX_-cells.

Overall, gene expression profiling along the trajectories confirmed that mild COVID-19 exhibits CD8^+^ T_RM_- and T_EX_-cells with good effector function, while in critical COVID-19 this effector function is drastically reduced possibly due to (persistent) inflammation-associated stress.

### Phenotypic heterogeneity of CD4^+^ T-cells in COVID-19 BAL

Amongst the 5 CD4^+^ T-cell clusters, we identified naïve CD4^+^ T-cells (T_N_), effector-memory T-cells (T_EM_), CD4^+^ T-helper-1 (T_H1_) cells, expressing high levels of immune-checkpoints (*HAVCR2*, *LAG3*, *PDCD1* and *CTLA4*), as well as CD4^+^ T-helper-17 (T_H17_) and CD4^+^ regulatory T-cells (TREG) (**Fig. 2b** for marker gene sets). Compared to non-COVID-19, we observed slightly less CD4^+^ T_H17_-cells, but more T_H1_-cells in COVID-19. Comparing mild *versus* critical COVID-19, we found T_H1_-cells to be significantly increased in the latter.

Slingshot revealed additional phenotypic heterogeneity, identifying central-memory CD4^+^ T_CM_-cells and stem cell-like memory CD4^+^ T_SCM_-cells (**Fig. 4a, b**), and constructed 3 trajectories, which were independently confirmed based on shared TCR clonotypes (**Fig. 4c**). Briefly, T_N_-cells connected closely with T_CM_-cells followed by T_EM_-cells, which branched-off into 3 different lineages to form T_H1_-cells, T_H17_-cells and T_SCM_-cells. Profiling of marker genes along these trajectories confirmed their functional annotation (**Fig. 4d**). Density plots stratified for T-cells from each subgroup revealed that COVID-19 was enriched for T-cells early and late in the T_H1_- and T_SCM_-trajectory, while *vice versa* T-cells from non-COVID-19 and normal lung were enriched halfway these trajectories (**Fig. 4e, f**). In the T_H17_-trajectory, COVID-19 BAL was strongly enriched for T-cells in the first half of the trajectory. Overall, CD4^+^ T-cells from mild COVID-19 behaved similarly as normal lung or non-COVID-19. Both TCR richness and evenness were reduced along the T_H1_-lineage from COVID-19, but not from non-COVID-19 (**Fig. 4g**). Notably, this reduction was most prominent in mild, but not in critical COVID-19 (**Fig. 4h**). In contrast, T_H17_-cells and T_SCM_-cells were characterized by a modest decrease in TCR richness only at the very end of their lineage, suggesting that mainly T_H1_-cells are selected for specific SARS-CoV-2 PAMPs/antigens.

**Fig. 4.**
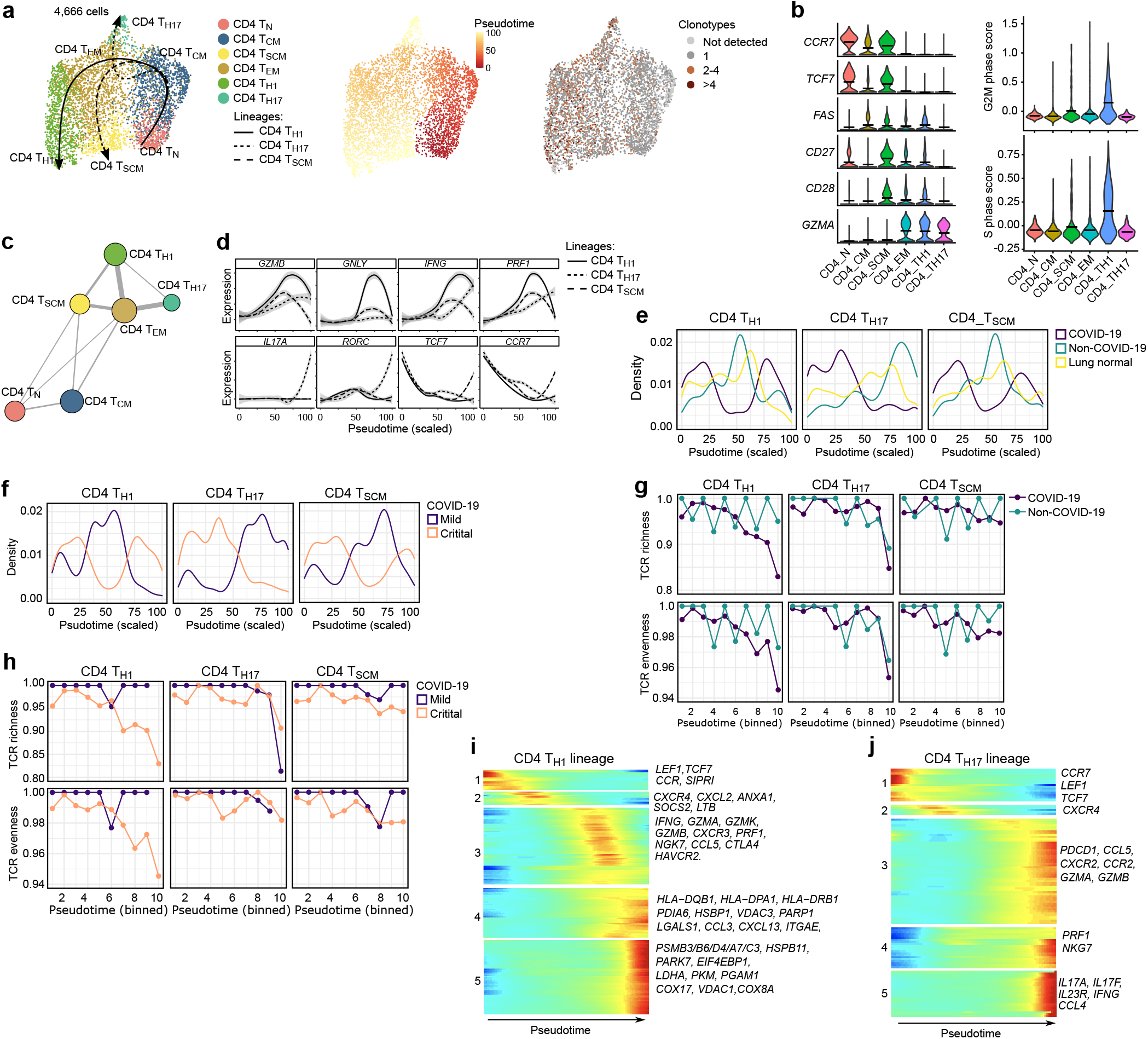
CD4^+^ T-cell developmental trajectories in mild and critical COVID-19 BAL. **a** UMAP with pseudotime trajectories based on Slingshot, showing 3 lineages (T_H1_-lineage, T_H17_-lineage and T_SCM_-lineage), color-coded for the CD4^+^ T-cell phenotypes (left), the pseudotime (middle) and the number of clonotypes (right). **b** Naïve and memory-related marker gene expression (left), and cell cycle scoring (right) reveal additional CD4^+^ T-cell subclusters. T_SCM_-cells are characterized by naïve marker genes (*CCR7*, *TCF7*), memory markers (*CD27*), cell proliferation but no *GZMA* expression. **c** Analysis of clonotype sharing (thickness indicates proportion of sharing) between the CD4^+^ T-cell subclusters. **d** Profiling of marker genes along these trajectories to confirm their functional annotation: *GZMB* and *IFNG* for the T_H1_-lineage, *IL17A* and *RORC* for the T_H17_-lineage, *TCF7* and *CCR7* for the T_SCM_-lineage, while *GNLY* and *PRF1* were plotted to highlighted T-cell effector function. **e** Density plots reflecting the number of T-cells along the 3 CD4^+^ T-cell lineages stratified for non-COVID-19, COVID-19 and normal lung. **f** Density plots reflecting the number of T-cells along the 3 CD4^+^ T-cell lineages stratified for mild *versus* critical COVID-19. **g-h** TCR richness and TCR evenness along the 3 CD4^+^ T-cell lineages comparing non-COVID-19 *versus* COVID-19 (**g**) and mild *versus* critical COVID-19 (**h**). **i-j** Gene expression dynamics along the CD4^+^ T_H1_-(**i**) and T_H17_-lineage (**j**). Genes cluster into 5 gene sets, each of them characterized by specific expression profiles, as depicted by a selection of marker genes characteristic for each set. Differences in trajectories were assessed by Mann-Whitney test. For CD4^+^ T_H1_ and CD4^+^ T_SCM_: COVID-19 *versus* non-COVID-19 and lung normal (P =1.4E-6 and 5.9E-37), For CD4^+^ T_Th17_: COVID-19 *versus* non-COVID-19 (P=9.7E-12), mild *versus* critical COVID-19 (P=1.3E-121).

Overall, this suggests that mild COVID-19 is characterized by more stable or differentiated T_H17_-cells’ activity, which is crucial for productive immunity against pathogens at mucosal surfaces^17^, whereas T_H1_-cells become entangled halfway in their trajectory. In critical COVID-19, T_H17_ cells completely fail to differentiate, while T_H1_-cells behave the opposite.

### Gene expression modelling along the CD4^+^ T_H1_- and T_H17_-lineage

Differential gene expression analysis along both lineages identified gene sets with specific expression profiles. In the T_H1_-lineage, the first gene set consisted of naïve (*LEF1, TCF7*) and undifferentiated (*CCR7*, *S1PR1*) T-cell markers (**Fig. 4i**; Supplementary information, **Table S5**). A second set was enriched for both pro- and anti-inflammatory markers (*CXCR4*, *CXCL2*, *ANXA1*, *SOCS2*, *LTB*), while a third set was characterized (halfway the trajectory) by an effector-like T_H1_-program based on expression of *IFNG*, granzymes (*GZMA*, *GZMK*, *GZMB*), *CXCR3*, *PRF1*, *NGK7*, *CCL5*, as well as *CTLA4* and *HAVCR2*. Finally, a fourth gene set was characterized by high HLA expression, auto-regulatory markers (*LGALS1*, *CCL3*), partial activation markers (*CXCL13*) and stress-response markers (*PDIA6*, *HSBP1*, *VDAC3*, *PARP1*) at the end of the trajectory, suggesting a complex mixture of a pro- and anti-inflammatory phenotype coupled with early-stress modulation. In a final fifth set, we noticed mitochondrial stress (*LDHA*, *PKM*, *COX17, VDAC1*, *COX8A*), an IL2 withdrawal-associated stressed phenotype (*MT1E*, *MT1X*), proteotoxic stress (*PSMB3/B6/D4/A7/C3*, *HSPB11*, *PARK7*, *EIF4EBP1*) and glycolysis (*PGAM1*) suggesting ‘terminal exhaustion’ at the end of the T_H1_-trajectory^18,19^. Overall, in mild COVID-19, the T_H1_-lineage was enriched for cells halfway the trajectory where expression of set 2-3 genes was most dominant, indicating increased T_H1_-effector function. In critical COVID-19, expression of sets 4-5 pre-dominated, suggesting inflammation-driven terminal exhaustion and severe dysregulation.

In the T_H17_-lineage, we also identified 5 gene sets (**Fig. 4j**; Supplementary information, **Table S6**): the first 2 sets with high expression early in the trajectory did not express markers indicative of T_H17_ function. Three other gene sets with high expression at the end of the trajectory were characterized by T-cell effector function (set 3: *PDCD1*, *CCL5*, *CXCR2*, *CCR2* and *GZMA/B*), expression of cytotoxic-activity genes (set 4: *NKG7* and *PRF1*) and T_H17_-cell associated interleukins (set 5: *IL17A*, *IL17F*, *IL23R,* as well as *IFNG* and *CCL4*). Notably, in mild COVID-19, cells were characterized by expression of genes belonging to set 3-5, while critical patients only expressed set 1-2 genes, completely failing to differentiate along the T_H17_ lineage.

Overall, this clearly indicates that mild COVID-19 is characterized by improved T_H1_- and T_H17_-effector functions that together mediate a highly controlled antiviral immune response, whose absence underlies critical COVID-19.

### Trajectory of monocyte-to-macrophage differentiation in COVID-19 BAL

In the 63,114 myeloid cells derived from BAL, we identified 9 phenotypes (**Fig. 5a**). Monocytes clustered separately from macrophages based on the absence of macrophage markers (*CD68*, *MSR1*, *MRC1*) and presence of monocyte markers (*IL1RN*, *FCN1*, *LILRA5*). Monocytes could be further divided into FCN1^high^, IL1B^high^ and HSPA6^high^ monocytes (**Fig. 5b**; Supplementary information, **Fig. S3a**), respectively, characterized by expression of classical monocyte markers (*IL1RN*, *S100A8/9*), pro-inflammatory cytokines (*IL1B*, *IL6*, *CCL3*, *CCL4*) and heat-shock proteins (*HSPA6*, *HSPA1A/B*). Based on *CSF1R, CSF3R* and *SPP1*, 3 monocyte-derived macrophage phenotypes could further be delineated, including CCL2^high^, CCL18^high^ and RGS1^high^ (Supplementary information, **Fig. S3b**). CCL2^high^ clusters were characterized by the pro-migratory cytokine *CCL2*, but also by several pro-(*CCL7*, *CXCL10*) and anti-inflammatory (*CCL13*, *CCL22*) genes, suggesting existence of an intermediate population of cells. In contrast, CCL18^high^ and RGS1^high^ cells expressed mainly anti-inflammatory genes (*CCL13*, *CCL18*, *PLD4, FOLR2*), as well as genes involved in receptor-mediated phagocytosis (*MERTK*, *AXL*). Finally, we identified MT1G^high^ macrophages (expressing numerous metallothioneins suggestive of oxidative stress or immune cell’s growth factor-withdrawal), a monocyte-derived (*FABP4*^*medium*^) and tissue-resident (*FABP4*^*high*^) alveolar macrophage cluster. The latter two populations were characterized by high expression of resident markers (*FABP4*, *PPARG*), anti-inflammatory (*CCL18*, *CCL22*) and antigen-presentation relevant MHC-I/II genes.

**Fig. 5.**
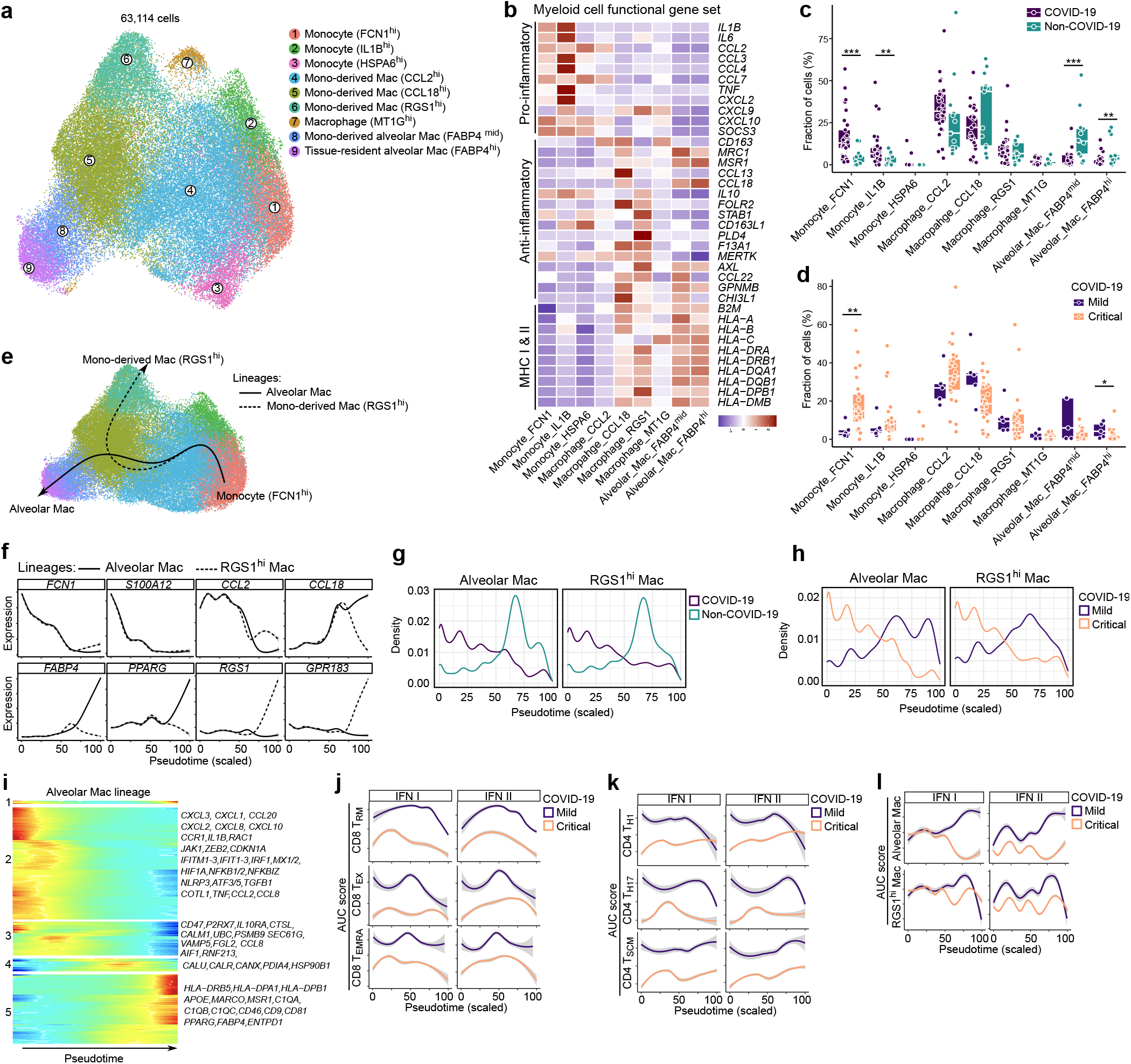
Monocyte-to-macrophage differentiation in COVID-19 BAL. **a** Subclustering of myeloid cells into 9 phenotypes, as indicated by the color-coded legend. **b** Heatmap showing myeloid cell phenotypes with corresponding functional gene sets. **c** Relative contribution of each cell type (in %) to COVID-19 *versus* non-COVID-19 BAL. **d** Relative contribution of each cell type (in %) to mild *versus* critical COVID-19 BAL. **e** Pseudotime trajectories for myeloid cells based on Slingshot, showing the common branch of FCN1^hi^ monocytes differentiating into either RGS1^hi^ monocyte-derived macrophages (RGS1^hi^-lineage) or FABP4^hi^ tissue-resident alveolar macrophages (alveolar lineage). **f** Profiling of marker genes along these trajectories to confirm their functional annotation: *FCN1, S100A12, CCL2, CCL18* for the common branch, *FABP4* and *PPARG* for the alveolar lineage, *RGS1* and *GPR183* for the RGS1-lineage. **g** Density plots reflecting the number of myeloid cells along the 2 lineages stratified for non-COVID-19 *versus* COVID-19. **h** Density plots reflecting the number of myeloid cells along the 2 lineages stratified for mild *versus* critical COVID-19. **i** Gene expression dynamics along the alveolar lineage. Genes cluster into 5 gene sets, each of them characterized by specific expression profiles, as depicted by a selection of genes characteristic for each cluster. **j-l** Profiling of IFN type I and II signalling along the 3 CD8^+^ (**j**) and CD4^+^ (**k**) T-cell lineages, and along the monocyte-macrophage lineage (l), comparing mild *versus* critical COVID-19. All P values were assessed by a Mann-Whitney test. * P<0.05, ** P<0.01, ***P<0.001. P values comparing COVID-19 *versus* non-COVID-19, and mild *versus* critical COVID-19 for density plots were all <10E-50.

We observed a significant increase in FCN1^high^ and IL1B^high^ monocytes in COVID-19 *versus* non-COVID-19, while *FABP4*^*medium*^ and *FABP4*^*high*^ alveolar macrophages were reduced (**Fig. 5c**). FCN1^high^ monocytes were significantly reduced in mild COVID-19, while alveolar macrophages were increased (**Fig. 5d**). Using Slingshot, we reconstructed two monocyte-to-macrophage lineages, consisting of a common branch of FCN1^high^ monocytes differentiating into IL1B^high^ monocytes, followed by CCL2^high^ and CCL18^high^ monocyte-derived macrophages. These subsequently branched into RGS1^high^ monocyte-derived macrophages (RGS1-lineage), or via *FABP4*^*medium*^ into *FABP4*^*high*^ tissue-resident macrophages (alveolar lineage; **Fig. 5e**). Monocyte marker expression decreased along both lineages, while macrophage marker expression increased (**Fig. 5f**). Density plots revealed that in COVID-19 cells were enriched in the first half of both lineages (**Fig. 5g**), confirming our above observations of monocyte enrichment in COVID-19. Comparing mild to critical COVID-19, we noticed that the former differentiated along both lineages, whereas in the latter monocytes completely failed to differentiate (**Fig. 5h**).

Modelling gene expression along the alveolar lineage revealed 5 gene sets (**Fig. 5i**; Supplementary information, **Table S7**). Sets 1 and 2 were characterized by inflammatory markers (*CXCL1-3*, *CCL20*, *CXCL8*, *CXCL10*, *CCR1*, *IL1B*), survival factors (*RAC1*, *JAK1*, *ZEB2*, *CDKN1A*), IFN-induced (anti-viral) genes (*IFITM1-3*, *IFIT1-3*, *IRF1*, *MX1/2*), hypoxia (*HIF1A*) and NF-κB (*NFKB1/2*, *NFKBIZ*) signalling early in the lineage, suggesting these monocytes to be characterized by a hyperinflammatory state, in which they prioritized inflammation rather than committing toward differentiation into macrophages. The third gene set was characterized by a possible CD47-based macrophage-suppressive phenotype, potentially aimed at dysregulating macrophage-activation (since CD47 is a well-established ‘don’t eat me’ signal striving to avoid auto-immunity)^20,21^. Moreover, based on expression of purinergic signalling (*P2RX7*), inflammasome or IL1-modulating factors (*NLRP3*, *IL1B*, *IL10RA*, *CTSL*, *CALM1*, *NFKB1*), endoplasmic reticulum (ER) stress capable of enabling ATP secretion (*UBC*, *PSMB9*, *SEC61G*, *ATF5*, *ATF3*), unconventional trafficking (*VAMP5*), fibrosis-related factors (*FGL2*, *TGFB1*, *COTL1*) and vascular inflammation (*TNF*, *AIF1*, *RNF213*, *CCL2*, *CCL8*) across sets 2 and 3 of these monocytes, we strongly suspect presence of extracellular ATP-driven purinergic-inflammasome signalling; especially given the high likelihood of extracellular ATP release from damaged epithelium in the context of acute viral infection. Importantly, this ATP-driven purinergic-inflammasome signalling pathway is a danger signalling cascade, which has been shown to facilitate ARDS-associated lung fibrosis and thus acts disease-worsening in this context^22–24^. Finally, set 4 and 5 genes were expressed at the end of the trajectory. Set 4 was characterized by expression of chaperone-coding genes (*CALU*, *CALR*, *CANX*, *PDIA4, HSP90B1*), which are crucial for robust functioning of the antigen-loading machinery for MHC molecules, whereas in set 5 there were clear signs of antigen presentation (expression of numerous MHC class II genes). Furthermore, set 5 comprised genes involved in receptor-mediated phagocytosis and post-phagocytic lipid degradation/metabolism: *APOE* for lipid metabolism, scavenger receptors *MARCO* and *MSR1*, complement activation (*C1QA*, *C1QB*, *C1QC* and *CD46*; that can also facilitate phagocytosis), viral infection-relevant inflammatory orientation (*CD81*, *CD9*), as well as anti-inflammatory markers (*PPARG*, *FABP4*)^25,26^. Similar gene sets were observed for the RGS1-lineage (Supplementary information, **Fig. S3c**), except for gene set 5, which exhibited expression of genes involved in chemokine signalling desensitization (*RGS1*), phagocytosis (*AXL*) and ATP clearance (*ENTPD1*)^27^.

Overall, this indicates that mild COVID-19 is characterized by functional pro-phagocytic and antigen-presentation facilitating functions in myeloid cells, whereas critical COVID-19 is characterized by disease-worsening characteristics related to monocyte-based macrophage suppression and ATP-purinergic signalling-inflammasome that may enable COVID-19 associated fibrosis and can worsen patient prognosis.

### Qualitative assessment of T-cell and monocyte/macrophage function in COVID-19

Next, although pseudotime inference usually allocates cells with a similar expression to the same pseudotime on the trajectory, we explored specific differences in gene expression along the pseudotime. We scored each cell using REACTOME pathway signatures and when comparing COVID-19 *versus* non-COVID-19 BAL, we observed consistently decreased IFN-signalling in non-COVID-19 T-cell and myeloid lineages (Supplementary information, **Fig. S4a**). In mild *versus* critical COVID-19, we observed that amongst several other pathways, IFN-(type I and II), interleukin (e.g., IL12 and IL6) and oligoadenylate synthetase (OAS) antiviral response signalling was increased in CD8^+^ T_RM_- and T_EX_-lineages (**Fig. 5j**; Supplementary information, **Fig. S4b-e**). The CD4^+^ T_H1_-lineage was similarly characterized by increased IFN-(type I and II) and interleukin (IL6, IL12, IL21) signalling in mild COVID-19 (**Fig. 5k**; Supplementary information, **Fig. S4f, g**).

Additionally, TRAF6-induced NF-κB and IRF7 activation, as well as TGFBR complex activation were increased. Similar effects were observed in the T_H17_-lineage (Supplementary information, **Fig. S4h, i**). The alveolar macrophage lineage was characterized by increased phagocytosis-related pathways (scavenging receptors, synthesis of lipoxins or leukotrienes) and IFN-signalling in mild COVID-19 (**Fig. 5l**; Supplementary information, **Fig. S4j-m**). *Vice versa*, IL10-signalling (which inhibits the IFN-response), chemokine receptor binding and ATF4-mediated ER stress response were increased in critical COVID-19.

Overall, while our trajectory and cell density analyses already indicated quantitative shifts in various cellular phenotypes comparing mild *versus* critical COVID-19, we noticed that also qualitatively immune cells from critical COVID-19 were severely dysfunctional.

### scRNA-seq of neutrophils, DCs and B-cells in COVID-19

We retrieved 14,154 neutrophils, which were subclustered into 5 phenotypes (**Fig. 6a, b**; Supplementary information, **Fig. S5a**). A first cluster consisted of ‘progenitor’ neutrophils based on *CXCR4* and *CD63*, and was also characterized by expression of the angiogenic factor *VEGFA* and cathepsins (*CTSA*, *CTSD*) (**Fig. 6c**). A second cluster consisted of few ‘immature’ neutrophils expressing *LTF*, *LCN2*, *MMP8*/9, *PADI4* and *ARG1*. Cluster 3 and 4 consisted of ‘inflammatory mature’ neutrophils, both expressing a signature footprint that highlights anti-pathogenic orientation of neutrophils^28^: cluster 3 expressed IFN-induced genes and calgranulins (*S100A8/9*, *S100A9* and *S100A12*), which can modulate inflammation, while cluster 4 expressed high levels of cytokines (*IL1B*) and chemokines (*CXCL8, CCL3*, *CCL4*). A final subset was characterized as ‘hybrid’ neutrophils due to their macrophage-like characteristics, i.e., expression of MHC class II and complement activation genes (*C1QB*, *C1QC, CD74*), cathepsins (*CTSB*, *CTSL*) and *APOE*. All neutrophil subclusters were more frequent in COVID-19 than non-COVID-19, but most significant changes were noticed for the ‘progenitor’ and ‘inflammatory mature’ neutrophils (**Fig. 6d**). Similar trends were observed in mild *versus* critical COVID-19, albeit non-significantly (**Fig. 6e**).

**Fig. 6.**
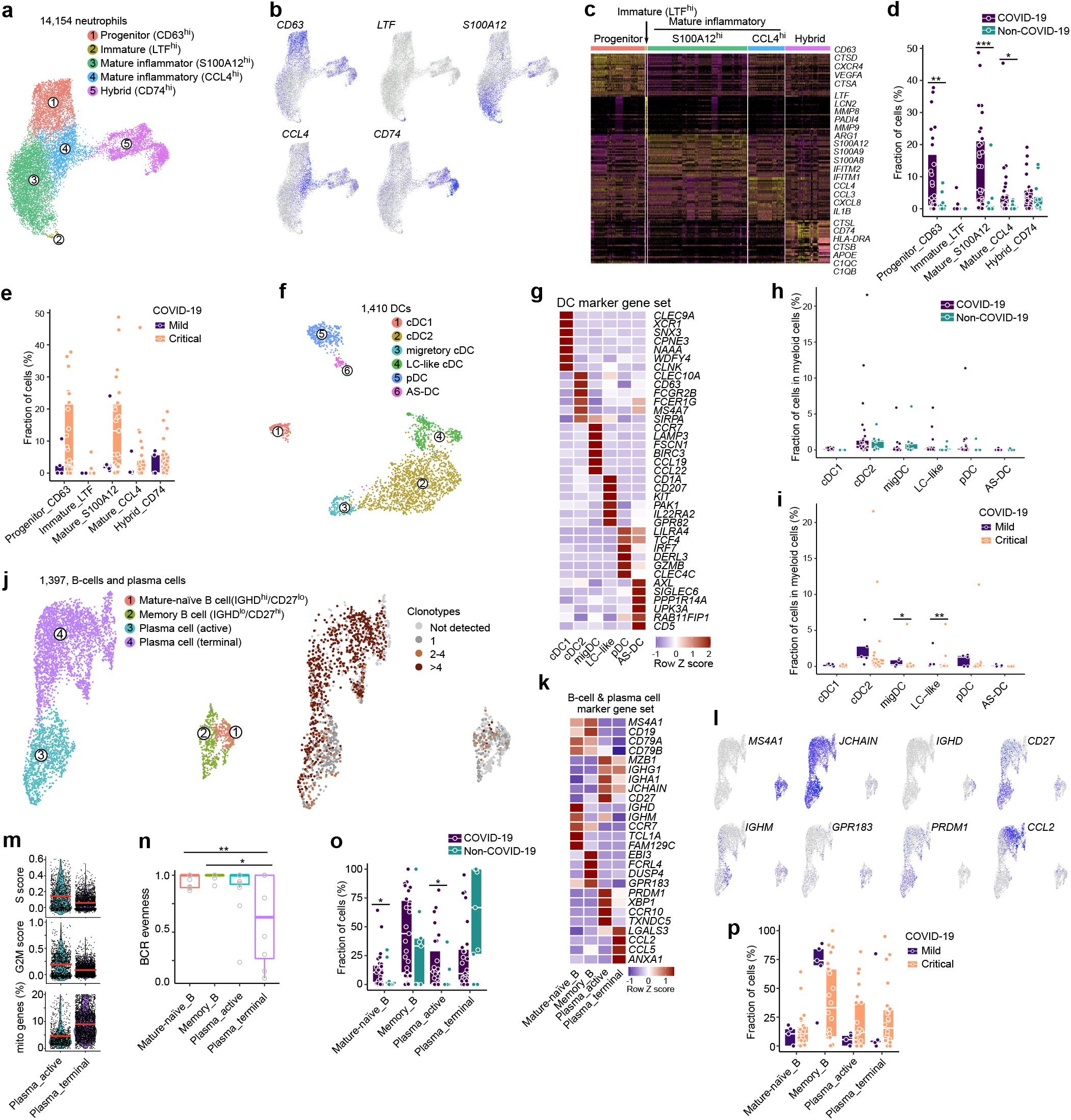
Neutrophil, dendritic cell and B-cell phenotypes in COVID-19 BAL. **a** Subclustering of neutrophils into 5 phenotypes, as indicated by the color-coded legend. **b** UMAP showing expression of a marker gene for each neutrophil phenotype. **c** Heatmap showing neutrophil phenotypes with corresponding marker genes and functional gene sets. **d** Relative contribution of each neutrophil phenotype (in %) to COVID-19 *versus* non-COVID-19. **e** Relative contribution of each neutrophil phenotype (in %) to mild *versus* critical COVID-19. **f** Subclustering of DC into 6 phenotypes, as indicated by the color-coded legend. **g** Heatmap showing DC phenotypes with corresponding marker genes and functional gene sets. **h** Relative contribution of each DC phenotype (in %) to COVID-19 *versus* non-COVID-19. **i** Relative contribution of each DC phenotype (in %) to mild *versus* critical COVID-19. **j** Subclustering of B-cells and plasma cells into 4 phenotypes, as indicated by the color-coded legend. **k** Heatmap showing B-cell and plasma cell phenotypes with corresponding marker genes and functional gene sets. **l** Feature plots of marker gene expression for each B-cell and plasma cell subcluster. **m** Violin plots showing cell cycle scores and mitochondrial gene expression by plasma cell subcluster. **n** B-cell receptor evenness in B-cell and plasma cell subclusters. **o** Relative contribution of each B-cell and plasma cell phenotype (in %) to COVID-19 versus non-COVID-19. **p** Relative contribution of each B-cell and plasma cell phenotype (in %) to mild versus critical COVID-19. P values were assessed by a Mann-Whitney test. * P<0.05, ** P<0.01, ***P<0.001.

We also identified 1,410 dendritic cells (DCs), which we could subcluster into 6 established populations (**Fig. 6f, g**; Supplementary information, **Fig. S5b, c**). None of these differed significantly between COVID-19 and non-COVID-19, while migratory DCs and Langerhans-cell-like DC were more frequent in mild *versus* critical COVID-19 (**Fig. 6h, i**).

Within the 1,397 B-cells, we obtained 4 separate clusters (**Fig. 6j**; Supplementary information, **Fig. S5d**). Follicular B-cells were composed of mature-naïve (CD27^−^) and memory (CD27^+^) B-cells. The former were characterized by a unique CD27^−^/IGHD^+^(IgD)/IGHM^+^(IgM) signature and give rise to the latter by migrating through the germinal center to form CD27^+^/IGHD^−^(IgD)/IGHM^−^(IgM) memory B-cells (**Fig. 6k, l**). Memory B-cells then further differentiate into antibody-secreting plasma cells (*IGHA1*, *IGHG1*, *JCHAIN*). A first cluster of ‘active’ plasma cells expressed high levels of *PRDM1*(Blimp-1) and *XBP1*, indicating high antibody-secretion capacity, while the latter was enriched for *CLL2* and *CCL5*, but also characterized by a reduced G2M and S cell cycle score and increased expression of mitochondrial genes, indicating ongoing stress (**Fig. 6m**). Notably, this population of ‘terminal’ plasma cells was also characterized by increased BCR clonality and reduced BCR evenness (**Fig. 6n**). Compared to non-COVID-19, mature-naïve B-cells and active plasma cells were increased in COVID-19, while terminal B-cells were reduced in CoVID-19, albeit non-significantly (**Fig. 6o**). There were no significant differences between mild *versus* critical COVID-19 (**Fig. 6p**). Overall, this suggests terminal B-cells in COVID-19 to be characterized by sub-optimal differentiation or activation, which may cause defective or counter-productive (possibly low-quality) antibody responses in COVID-19.

### SARS-CoV-2 viral particles in epithelial and immune cells

Finally, we retrieved 22,215 epithelial cells, which we subclustered into 7 distinct clusters (**Fig. 7a, b**; Supplementary information, **Fig. S5e, f**), the largest 3 clusters consisting of secretory, ciliated and hillock lung epithelial cells. The basal population (*KRT5*, *AQP3* and *SPARCL1*), representing stem cell epithelial cells responsible for epithelial remodelling upon lung injury, was significantly enriched in COVID-19 *versus* non-COVID-19, as well as ionocytes, which is another rare epithelial cell type that regulates salt balance (**Fig. 7c**). There were no significant differences between mild *versus* critical COVID-19 (**Fig. 7d**). Interestingly, *ACE2* and *TMPRSS2* expression was increased in COVID-19 *versus* non-COVID-19, with 21% and 2.3% of epithelial cells being positive, respectively (**Fig. 7e, f**). We then assessed in which cells we retrieved sequencing reads mapping to the SARS-CoV-2 genome, identifying 3,773 positive cells from 17 out of 31 COVID-19 patients. Surprisingly, this revealed a higher overall number of reads mapping to lymphoid and myeloid than epithelial cells (**Fig. 7g**). Stratification for each of the 11 SARS-CoV-2 open-reading frames (ORF) using Viral-Track revealed that the RNA encoding for spike protein (*S*), which interacts which ACE2 during viral entry of the cell, was almost exclusively detected in epithelial cells, which were also the only cells expressing *ACE2* and *TMPRSS2* (**Fig. 7g**). In contrast, the nucleocapsid protein (*N*), and to a lesser extent the ORF10 and ORF1a-encoding mRNAs were detected in myeloid and lymphoid cells at much higher levels than in epithelial cells (**Fig. 7h**). Further stratification into cell types revealed that neutrophils and to a limited extent also monocytes, contained most reads mapping to *N* (**Fig. 7i**). This might suggest that neutrophils are the main cell type interacting with SARS-CoV-2 viral particles/infected cells and account for the highest procurement of viral material, in line with their role as first innate immune responders to infection^29^. Differential gene expression of *N*-positive *versus N*-negative neutrophils identified upregulation of transcription factor *BCL6*, which promotes neutrophils survival and inflammatory response following virus infection, and numerous IFN-induced genes (*IFITM3*, *IFIT1-3*, *MX1/2*, ISG15, *RSAD2*; **Fig. 7j**)^30^. No such enrichment was observed in monocytes nor macrophages (Supplementary information, **Fig. S5g-i**). Pathway analysis on differentially-expressed genes revealed IFN-signalling using REACTOME and Response_to_virus using GO for genes upregulated in *N*-containing neutrophils (**Fig. 7k, l**). Notably, amongst the different neutrophil phenotypes, *N* was most strongly enriched in ‘inflammatory mature’ neutrophils expressing calgranulins (**Fig. 7m**). As expected, significantly more *N* was present in critical *versus* mild COVID-19^31^.

**Fig. 7.**
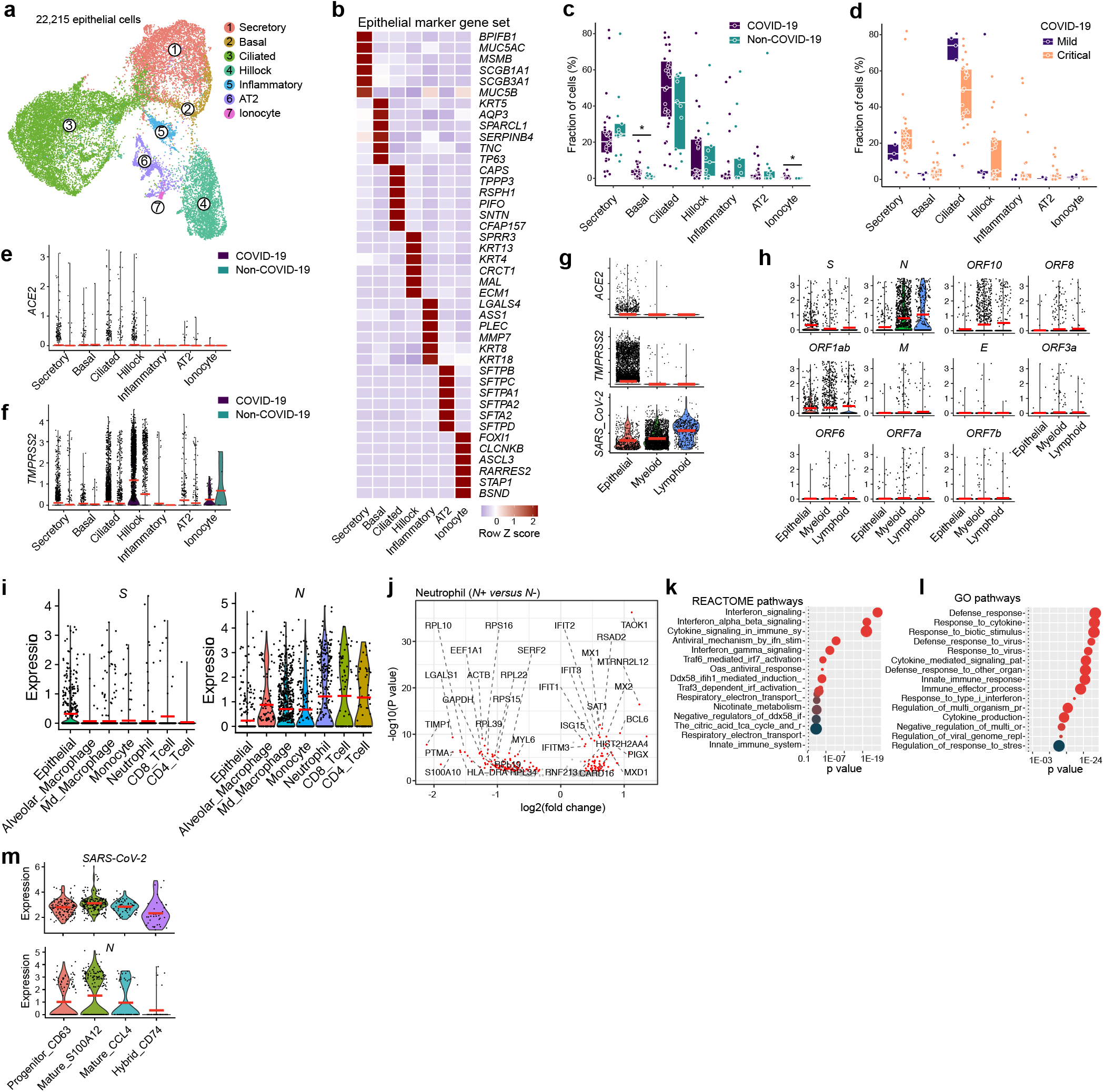
SARS-CoV-2 RNA detection in epithelial and immune cells. **a** Subclustering of epithelial cells into 7 phenotypes, as indicated by the color-coded legend. **b** Heatmap showing epithelial cell phenotypes with corresponding marker genes. **c** Relative contribution of each epithelial cell phenotype (in %) to COVID-19 *versus* non-COVID-19. **d** Relative contribution of each epithelial cell phenotype (in %) to mild *versus* critical COVID-19. **e-f** Expression level of *ACE2* (**e**) and *TMPRSS2* (**f**) by epithelial cell subclusters, comparing COVID-19 *versus* non-COVID-19. **g** Expression levels of *ACE2*, *TMPRSS2* and *SARS-CoV-2* (cells with viral reads) RNA in epithelial, myeloid and lymphoid cells from COVID-19. **h** Detection of 11 SARS-CoV-2 open-reading frames in epithelial, myeloid and lymphoid cells from COVID-19. **i** Detection of spike protein (*S*) and nucleocapsid protein (*N*) encoding viral RNA in epithelial cells and immune cell subclusters from COVID-19. Cell types with <50 positive cells are not shown. **j** Differential gene expression of *N*-positive *versus N*-negative neutrophils from 17 COVID-19 patients in which viral reads were detected. **k-l** REACTOME (**k**) and GO (**l**) pathway analysis on IFN-signalling and response-to-virus signalling, comparing *N*-positive *versus N*-negative neutrophils from 17 COVID-19 patients in which viral reads were detected. **m** Detection of reads mapping to *SARS-CoV-2* and to *N* in neutrophil subclusters from COVID-19 BAL. P values were assessed by a Mann-Whitney test. * P<0.05, ** P<0.01, ***P<0.001.

### Cell-to-cell communication to unravel the immune context of COVID-19 BAL

Since, our data on the one hand reveal that neutrophils were involved in cleaning up viral particles/virus-infected cells, yet T-cell and monocyte-to-macrophage lineages were significantly disrupted in critical COVID-19, we explored the (predicted) interactome between these cell types to gain more refined insights. First, we calculated interactions between cell types (P≤0.05) separately for mild and critical COVID-19, then we assessed differences in the number of specific interactions. Neutrophils were characterized by a low number of specific interactions that were slightly more frequent in critical *versus* mild COVID-19. *Vice versa*, numerous specific interactions were predicted between all other immune and epithelial cells, especially in mild COVID-19 (**Fig. 8a, b**; Supplementary information, **Fig. S6**).

**Fig. 8.**
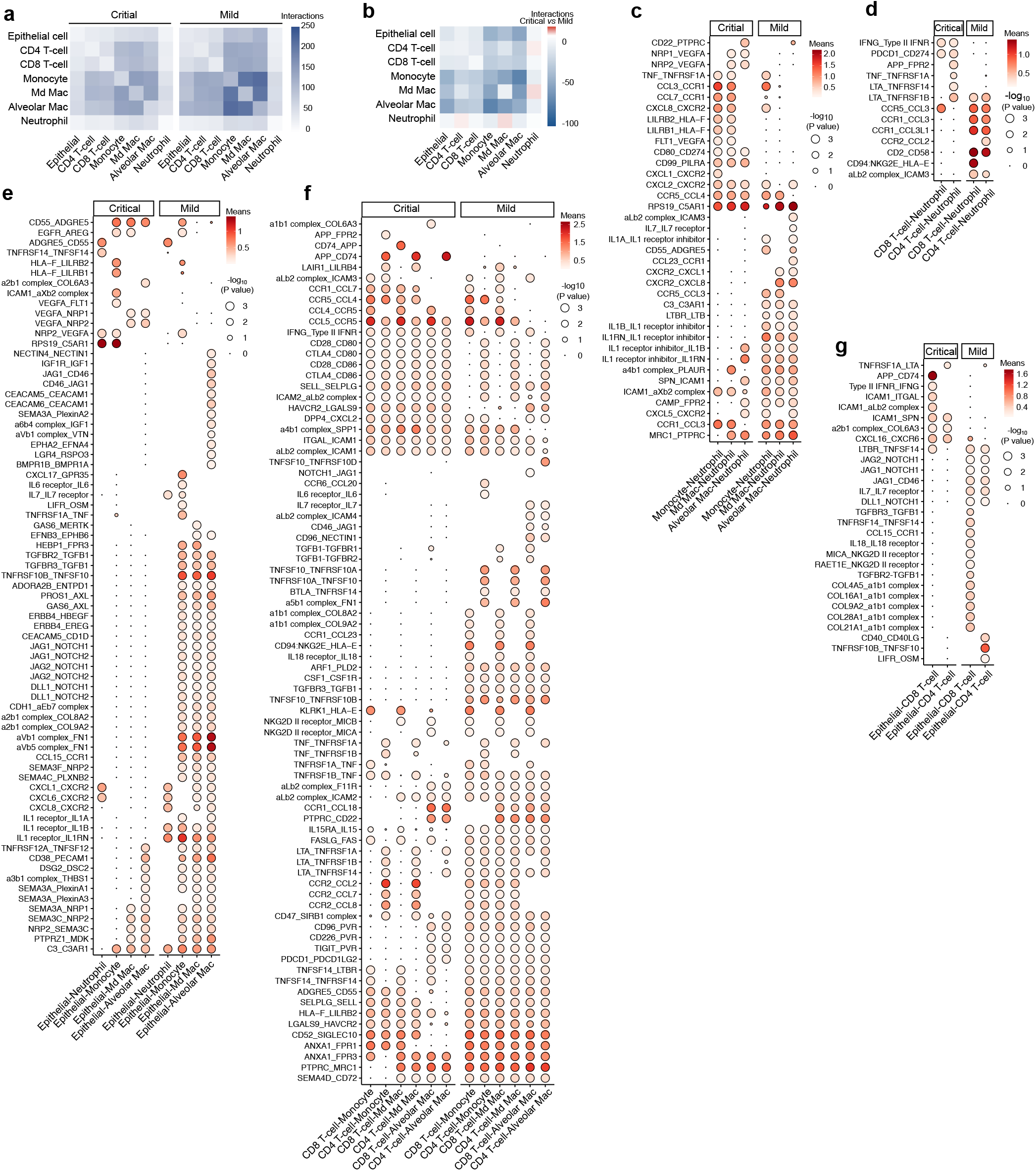
Cell-to-cell communication between epithelial and immune cells. **a** Number of predicted interactions (P≤0.05) between monocytes, macrophages, T-cells, neutrophils and epithelial cells based on CellPhoneDB in critical (left panel) and mild (right panel) COVID-19. **b** Differences in the number of predicted interactions, comparing mild *versus* critical COVID-19, showing generally more interactions in mild COVID-19. **c** Predicted interactions between monocytes/macrophages and neutrophils, comparing critical *versus* mild COVID-19. **d** Predicted interactions between T-cells and neutrophils, comparing critical *versus* mild COVID-19. **e** Predicted interactions between epithelial and myeloid cells, comparing critical *versus* mild COVID-19. **f** Predicted interactions between T-cells and monocytes/macrophages, comparing critical *versus* mild COVID-19. **g** Predicted interactions between T-cells and epithelial cells, comparing critical *versus* mild COVID-19.

In critical COVID-19, specific interactions between monocytes/macrophages and neutrophils almost always involved pro-migratory interactions (*FLT1*, *NRP1* or *NRP2*/*VEGFA*, *CXCL1* or *CXCL2* or *CXCL8*/*CXCR2*, *CCL3* or *CCL7*/*CCR1*), coupled with immune-inhibitory interactions, such as *LILRB1* or *LILRB2*/*HLA-F* and *RPS19*/*C5AR1*, which also induce neutrophil dysfunction (**Fig. 8c**)^32^. A few stimulatory T-cell to neutrophil interactions were observed, including *IFNG*/*type II IFNR*, *PDCD1*/*CD274*, *LTA*/*TNFRSF1A* or *TNFRSF1B* (**Fig. 8d**), while specific epithelial cell-to-neutrophil interactions were limited to a mixture of myeloid immunosuppression (*RPS19*/*C5AR1*) and viral infection-relevant pro-inflammatory signals (*TNFRSF14*/*TNFSF14*) (**Fig. 8e**). Amongst T-cell and monocytes/macrophages, some immune-stimulatory or auto-regulatory interactions were seen (*CTLA4* or *CD28*/*CD80* or *CD86*, *CCL5/CCR5*) (**Fig. 8f**), but specific epithelial to T-cell interactions in critical COVID-19 were limited to pro-inflammatory ICAM1-mediated interactions (**Fig. 8g**).

A very different scenario was observed in mild COVID-19 (**Fig. 8c-g**). Amongst the numerous interactions between monocytes/macrophages and neutrophils, we noticed interleukin signalling (bi-directional *IL1B*, *IL1A, IL1RN/A* signalling, *IL7*/*IL7R*, *CXCR2*/*CXCL1* or *CXCL8*), but also *MRC1*/*PTPR* (phagocytosis) and *LTBR/LTB* (pro-inflammation). Between T-cells and neutrophils specific interactions involved *CCR1*/*CCL3* or *CCL3L1* (pro-inflammation), *CD2*/*CD58* (co-stimulatory/immunogenic pathway) and *CD94:NKG2E*/*HLA-F* (anti-viral immune-surveillance), whereas between epithelial cells and neutrophils, *IL1R*/*IL1A* or *IL1B* or *IL1R* interactions were most pronounced (which can facilitate productive neutrophil immunity in an immune-controlled/immunogenic context)^28,33,34^. Numerous interactions were also observed between epithelial cells and monocyte/macrophages: *GAS6* or *PROS1*/*AXL* (receptor-mediated phagocytosis), *ADORA2B*/*ENTPD1* (extracellular ATP degradation/suppression), *CD83*/*PECAM1* (immune activation) and semaphorins interacting with their plexin and NRP receptors (tissue re-modelling and repair). Between epithelial cells and T-cells, we observed mainly co-stimulatory (*CD46*/*JAG1*, *CD40LG*/*CD40*, *IL7R*/*IL7*, *MICA* or *RAET11/NKG2D receptor*) and tissue repair interactions (*TGFB1/TGFR2* and *TGFB1/TGFBR3)*, while amongst T-cells and monocytes/macrophages, there were amongst others, co-stimulatory (*LTA*/*TNFRSF1A* or *TNFRSF1B* or *TNFRSF14*, *TNFSF10*/*TNFRSF10B*) or tissue-repair factors (*CSF1*/*CSFR1*, *TGFBR3*/*TGFB1*, *IL15RA*/*IL15*), mediators of T-cell homeostasis and cytotoxicity (*FASLG*/*FAS*) and antiviral immune surveillance (*NKG2D II receptor*/*MICB* or *MICA*).

## DISCUSSION

Based on scRNA-seq data obtained from BAL fluid, we were able to perform deep-immune profiling of the adaptive and innate immune cell landscape within the main locale of COVID-19 pathology. A particular strength of our study is the profiling of BAL from a fairly large cohort of COVID-19 patients (n=31), enabling statistically meaningful and robust comparisons between mild and critical disease severity subgroups (in contrast to initial COVID-19 publications that profiled <10 patients)^7^. Importantly, our control group also consisted of non-COVID-19 pneumonia cases (n=13), instead of healthy controls. Since the latter are likely to differ on almost every immunological parameter-level relative to COVID-19, our strategy enhances qualitative clarity of immunological conclusions. Finally, due to the fact that we profiled >116,000 single-cells, we could infer pseudotime trajectories for both T-cells and myeloid cells. Such method is particularly attractive since it allows modelling of gene expression changes along the inferred trajectories, thereby generating data at a much greater resolution. Overall, this allowed us to draw the following key conclusions regarding what distinguishes a critical from a mild COVID-19 disease course:

Firstly, CD8^+^ T-cells exhibited good effector functions along their resident-memory and partially-exhausted lineages in mild COVID-19, while also CD4^+^ T-cells showed increased effector or disease-resolving functions in T_H1_- and T_H17_-lineages. In critical COVID-19, T-cells were highly dysregulated, either failing to differentiate (T_17_- and T_RM_-lineage) or exhausting excessively, thereby leading to metabolic disparities, dysregulation of their immunological interface with myeloid cells and/or a dysregulated chronic hyper-inflammatory phenotype (T_H1_ and T_EX_-lineage). Notably, we observed that mild *versus* critical COVID-19, not only differed quantitatively in terms of the number of T-cells exhibiting a good T-cell effector function, but also qualitatively, in terms of consistently lower activation levels of the type 1 and II IFN (anti-viral) signalling pathways (amongst several other pathways). Overall, this showed that T-cells in mild COVID-19, unlike those in critical COVID-19, were cross-talking better with their lung-localised microenvironment thereby facilitating ‘ordered’ immune reactions capable of resolving, rather than exacerbating, inflammation and tissue repair ^35^.

Secondly, in mild COVID-19 monocytes exhibited a pronounced pro-inflammatory phenotype, but then differentiated into macrophages characterized by anti-inflammatory, pro-phagocytic and antigen-presentation facilitating functions. This suggests that in these patients, macrophages might be immunologically ‘silently’ cleaning the dying/dead epithelial cells (as well as other immune cells meeting their demise due to inflammation), hence contributing to degradation and dilution of the viral load in COVID-19 BAL. Such pro-homeostatic activity of macrophages is well-established to aid in disease amelioration and inflammation resolution^36^. In critical COVID-19, monocytes were instead characterized by a chronically hyper-inflamed phenotype with characteristics of an ATP-inflammasome-purinergic signalling-based fibrosis, which can promote worse disease outcome by contributing to development of ARDS. This danger signalling pathway is hypothesized to be part of the chronology of events during SARS-CoV-2 infection, but its genetic footprints have not been documented as we report here^37^. Considering that fully-differentiated macrophages are much more efficient in clearing large debris or cellular corpses (e.g. infected dead/dying lung epithelia or dead neutrophils) than monocytes or neutrophils, their dysfunction in critical COVID-19 may explain the excessive accumulation of lung epithelial (as well as dead immune cell) debris and alveolar dyshomeostasis coupled with dysregulated coagulopathy^38–42^.

Lastly, based on the presence of sequencing reads mapping to the gene encoding for viral protein S, which is needed to infect cells via ACE2 and TMPRSS2 receptors, we propose that SARS-CoV-2 largely infects epithelial cells (as primary targets of excessive pathological replication and propagation), but not necessarily lymphoid or myeloid cells (although we cannot exclude yet that some virions might be capable of ‘latently’ entering these cells without showing pathological replication or propagation). Interestingly, we also detected reads mapping to the nucleocapsid protein (*N*) encoding gene mainly in neutrophils, but to some extent also in other lymphoid or myeloid cells, especially monocytes. This suggests that neutrophils might be heavily involved in viral clearance of SARS-CoV-2 – as is the case in most viral pathologies^29^. Indeed, we observed that ‘inflammatory mature’ neutrophils, which exhibited an anti-pathogenic orientation with pronounced degranulating activity, contained most of the viral *N* sequences (amongst all other neutrophil phenotypes). Moreover, *N*-positive neutrophils exhibited increased expression of IFN-induced (anti-viral) genes, compared to *N*-negative neutrophils. Some sequencing reads also mapped to *ORF10* and *ORF1ab*, but not the other viral protein-encoding genes. We suspect this is due to increased stability of *N*, *ORF10* and *ORF1ab* RNA compared to other viral ORFs. In conclusion, these data suggest that the neutrophil’s positioning in an immune-inhibitory (adverse) environment, with disrupted T-cell effector/regulatory function as well as mostly inhibitory or dysregulated interactions with other (myeloid) immune cells, might explain their failure in controlling disease progression, thereby leading to critical COVID-19 pathology.

Our findings bear important therapeutic relevance. The RECOVERY trial recently claimed that dexamethasone reduces death by one-third in hospitalised patients with critical COVID-19 (unpublished data). Dexamethasone has indeed been shown to dampen myeloid inflammatory signalling (notably IL-1 and IL-6 release), reduce neutrophil inflammation^43^, promote an ‘M2-like” macrophage phenotype, which has anti-inflammatory and phagocytic traits^44^, as well as to maintain clonal balance in T-cells^45^. Given the findings we report here, the therapeutic effects of dexamethasone are not entirely unexpected. Our data also suggest that neutrophils are key players in the acute phase of the infection. However, prolonged neutrophil inflammation might also cause excessive collateral lung damage and be detrimental to the host, as suggested by autopsy reports^46^. In this regard, the immunomodulatory antibiotic azithromycin might be a promising therapy for COVID-19 when administered early in the disease course. Acutely administered azithromycin enhances degranulation and the oxidative burst by neutrophils in response to a stimulus, yet this is followed by a subsequent decrease of oxidative burst capacity and increase in neutrophil apoptosis^47^. We are therefore eagerly awaiting results from large-scale randomised trials with azithromycin for COVID-19.

Nevertheless, there are also limitations to our study. For instance, we observed evidence of counter-productive (possibly low-quality) antibody response-related signatures in COVID-19, but failed to perform an in-depth study in this area. Additional studies performing scRNA- and scBCR-seq on serially-collected samples during disease are needed to reinforce this observation. Also, several COVID-19 patients were treated with the antiviral drugs remdesivir, which targets the viral RNA-dependent RNA polymerase, or hydroxychloroquine, which has immunomodulatory traits and is still controversial with respect to its therapeutic effects on disease outcome^48–50^. Of note, we did not detect major patient-specific cell clusters nor other type of outliers during our analyses.

In conclusion, we used single-cell transcriptomics to characterize the innate and adaptive lung immune response to SARS-CoV-2. We observed marked changes in the immune cell compositions, phenotypes as well as immune cross-talks during SARS-CoV-2 infection and identified several distinguishing immunological features of mild *versus* critical COVID-19. We also documented genetic footprints of several crucial immunological pathways that have been extensively hypothesized, but not always systematically confirmed, to be associated with COVID-19 pathology and SARS-CoV-2 infection biology. We believe that this work represents a major resource for understanding lung-localised immunity during COVID-19 and holds great promise for the study of COVID-19 immunology, immune-monitoring of COVID-19 patients and relevant therapeutic development.

## MATERIALS AND METHODS

### Patient cohort, sampling and data collection

22 COVID-19 patients and 13 non-COVID-19 pneumonitis patients in this study were enrolled from the University Hospitals Leuven, between March 31^th^ 2020 and May 4^th^ 2020. Disease severity was defined as ‘mild’ or ‘critical’, based on the level of respiratory support at the time of sampling. Specifically, ‘mild’ patients required no respiratory support or supplemental oxygen through a nasal cannula, whereas ‘critically ill’ patients were mechanically ventilated or received extracorporeal membrane oxygenation.

The demographic and disease characteristics of the prospectively recruited patients studied by scRNA-seq are listed in Supplementary Table 1. Diagnosis of COVID-19 was based on clinical symptoms, chest imaging and SARS-CoV-2 RNA-positive testing (qRT–PCR) on a nasopharyngeal swab and/or BAL fluid sample. Non-COVID-19 pneumonitis cases all tested negative for SARS-CoV-2 RNA using a qRT-PCR assay on BAL.

All 35 patients underwent bronchoscopy with BAL as part of the standard of medical care, because of i) high clinical suspicion of COVID-19 yet negative SARS-CoV-2 qRT-PCR on nasopharyngeal swab ii) established COVID-19 with clinical deterioration, to rule out opportunistic (co-)infection and/or to remove mucus plugs. Lavage was performed instilling 20cc of sterile saline, with an approximate retrieval of 10cc. 2-3cc of the retrieved volume was used for clinical purposes. The remaining fraction was used for scRNA-seq.

The retrieved BAL volume was separated into two aliquots, as explained above, at the bedside by the performing endoscopist. The aliquot used for scRNA-seq was immediately put on ice and transported to a Biosafety Level 3 Laboratory (REGA Institute, KU Leuven) for scRNA-seq.

Demographic, clinical, treatment and outcome data from patient electronic medical records were obtained through a standardized research form in Research Electronic Data Capture Software (REDCAP, Vanderbilt University). This study was conducted according to the principles expressed in the Declaration of Helsinki. Ethical approval was obtained from the Research Ethics Committee of KU / UZ Leuven (S63881). All participants provided written informed consent for sample collection and subsequent analyses.

### scRNA-seq, scTCR-seq and scBCR-seq profiling

BAL fluid was centrifuged and the supernatant was frozen at −80°C for further experiments. The cellular fraction was resuspended in ice-cold PBS and samples were filtered using a 40µm nylon mesh (ThermoFisher Scientific). Following centrifugation, the supernatant was decanted and discarded, and the cell pellet was resuspended in red blood cell lysis buffer. Following a 5-min incubation at room temperature, samples were centrifuged and resuspended in PBS containing UltraPure BSA (AM2616, ThermoFisher Scientific) and filtered over Flowmi 40µm cell strainers (VWR) using wide-bore 1 ml low-retention filter tips (Mettler-Toledo). Next, 10 µl of this cell suspension was counted using an automated cell counter to determine the concentration of live cells. The entire procedure was completed in less than 1.5 h.

Single-cell TCR/BCR and 5’ gene expression sequencing data for the same set of cells were obtained from the single-cell suspension using the Chromium™ Single Cell 5’ library and Gel Bead & Multiplex Kit with the Single Cell V(D)J Solution from 10x Genomics according to the manufacturer’s instructions. Up to 5,000 cells were loaded on a 10x Genomics cartridge for each sample. Cell-barcoded 5’ gene expression libraries were sequenced on an Illumina NovaSeq6000, and mapped to the GRCh38 human reference genome using CellRanger (10x Genomics, v3.1). V(D)J enriched libraries were sequenced on an Illumina HiSeq4000 and TCR and BCR alignment and annotation was achieved with CellRanger VDJ (10x Genomics, v3.1).

### Single-cell gene expression analysis

Raw gene expression matrices generated per sample were merged and analysed with the Seurat package (v3.1.4)^51^. Cell matrices were filtered by removing cell barcodes with <301 UMIs, <151 expressed genes or >20% of reads mapping to mitochondrial RNA. We opted for a lenient filtering strategy to preserve the neutrophils, which are transcriptionally less active (lower transcripts and genes detected). The remaining cells were normalized and the 3000 most variable genes were selected to perform a PCA analysis after regression for confounding factors: number of UMIs, percentage of mitochondrial RNA, patient ID and cell cycle (S and G2M phase scores calculated by the CellCycleScoring function in Seurat), interferon response (BROWNE_INTERFERON_RE-SPONSIVE_GENES in the Molecular Signatures Database or MSigDB v6.2), sample dissociation-induced stress signatures^52^, hypoxia signature^53^. This PCA and graph-based clustering approach however resulted in some highly patient specific clusters, which prompted us to perform data integration using anchor-based CCA in Seurat (v3) package between patients to reduce the patient-specific bias. And this was performed after excluding cells from an erythrocyte cluster (primarily from a single patient) and a low-quality cell cluster. After data integration, 3000 most variable genes were calculated by FindVariableFeatures function, and all the mitochondrial, cell cycle, hypoxia, stress and interferon response genes (Pearson correlation coefficient > 0.1 against scores of the above-mentioned signatures calculated by AddModuleScore function in Seurat) were removed from the variable genes. In addition, we also removed common ambient RNA contaminant genes, including hemoglobin and immunoglobulin genes, as well as T-cell receptor (TRAVs, TRBVs, TRDVs, TRGVs) and B-cell receptor (IGLVs, IGKVs, IGHVs) genes, before downstream analyses.

### scRNA-seq clustering for cell type identification

For the clustering of all cell types, principal component analysis (PCA) was applied to the variable genes of dataset to reduce dimensionality. The selection of principal components was based on elbow and Jackstraw plots (usually 25-30). Clusters were calculated by the FindClusters function with a resolution between 0.2 and 2, and visualised using the Uniform Manifold Approximation and Projection for Dimension Reduction (UMAP) reduction. Differential gene-expression analysis was performed for clusters generated at various resolutions by both the Wilcoxon rank sum test and Model-based Analysis of Single-cell Transcriptomics (MAST) using the FindMarkers function^51^. A specific resolution was selected when known cell types were identified as a cluster at a given resolution, but not at a lower resolution with the minimal constraint that each cluster has at least 10 significantly differentially expressed genes (FDR <0.01, 2-fold difference in expression compared to all other clusters). Annotation of the resulting clusters to cell types was based on the expression of marker genes.

### Integration of publicly available datasets and identification of cell subtypes

We additionally processed scRNA-seq data on COVID-19 BAL fluid by Liao et al. and on normal lung samples by Reyfman et al. and Lambrechts et al. as described above^7,9,10^. The former two datasets were *de novo* clustered and annotated, and cell type annotation of the last dataset was used as previously described^11^. For cell subtype identification, the main cell types identified from multiple datasets were pooled, integrated, and further subclustered using the similar strategy, except that the constant immunoglobulin genes were not excluded for B-cell and plasma cell subclustering. Finally, doublet clusters were identified based on: 1) expression of marker genes from other cell (sub)clusters, 2) higher average UMIs as compared to other (subclusters), and 3) a higher than expected doublets rate (> 20%), as predicted by both DoubletFinder (v2)^54^ and Scrublet^55^ and the clustering was re-performed in the absence of the doublet clusters.

### Trajectory inference analysis

The R package Slingshot was used to explore pseudotime trajectories/potential lineages in T- and myeloid cells^56^. The analyses were performed for CD8^+^ and CD4^+^ cell phenotypes separately, with T_MAIT_-, T_γδ_- and T_REG_-cells excluded due to their unique developmental origin. For each analysis, PCA-based dimension reduction was performed with differentially expressed genes of each phenotype, followed by two-dimensional visualization with UMAP. Graph-based clustering (Louvain) identified additional heterogeneity for some phenotypes, as described in the manuscript for CD4^+^ T-cells. Next, this UMAP matrix was fed into SlingShot, with naïve T-cells as a root state for calculation of lineages and pseudotime. Similar approach was applied to the monocyte-macrophage differentiation trajectory inferences.

### Assessing the TCR and BCR repertoires

We only considered productive TCR/BCRs, which were assigned by the CellRanger VDJ pipeline. Relative clonotype richness^57^, defined as the number of unique TCRs/BCRs divided by the total number of cells with a unique TCR/BCR, was calculated to assess clonotype diversity. Relative clonotype evenness^58^, was defined as inverse Simpson index divided by species richness (number of unique clonotypes).

### Inflammatory pathways and gene set enrichment analysis and tradeSeq

The REACTOME pathway activity of individual cells was calculated by AUCell package (v1.2.4)^59^. And the differential activity between lineages along the trajectories were calculated using TradeSeq^60^. Pathways with median fold change >3 and an adjusted p-value < 0.01 were considered as significantly changed. GO and REACTOME geneset enrichment analysis were performed using hypeR package^61^; geneset over-representation was determined by hypergeometric test.

### SARS-CoV-2 viral sequence detection

Viral-Track was used to detect SARS-CoV-2 reads from BAL scRNA-seq data (reference genome NC_045512.2), as previously described^8^. The initial application was aimed to identify SARS-CoV-2 reads against thousands of other viruses, and thus the STAR indexes for read alignment were built by combining the human (GRCh38) genome reference with thousands of virus refence genomes from viruSITE. Since the likelihood of co-infection with multiple viruses (>2) is low in COVID-19 patients^8^, we adapted the Viral-Track pipeline to reduce computation time and increase sensitivity. Briefly, instead of directly processing raw fastq reads, we took advantage of BAM reads generated for scRNA-seq data, which mapped to human genome by the CellRanger pipeline as described above. The BAM files were filtered to only keep reads with cell barcodes annotated in the scRNA-seq analysis using subset-bam tools (10x Genomics). Then the corresponding unmapped BAM reads were extracted using samtools and converted to fastq files using bamtofastq tool to be further processed by UMI-tools for cell barcode assignment before feeding into Viral-Track pipeline. These unmapped reads, which contain potential viral sequences, were aligned using STAR to SARS-CoV-2 reference genome, with less stringent mapping parameter (outFilterMatchNmin 25-30), as compared to the original Virial-Track pipeline. Our approach identified 17 SARS-CoV-2 positive patients from a total of 31 COVID-19 patients, including 3 patients that were previously not detected using original Viral-Track pipeline by Bost et al. None of the patients among the 13 non-COVID-19 patients were detected as SARS-CoV-2 positive, suggesting our adapted pipeline does not result in major false-positive detection. For the detection of 11 SARS-CoV-2 ORFs or genes, a GTF annotation file was generated according to NC_045512.2^62^ for counts matrix using Viral-Track. The viral gene counts of each barcoded cells were integrated into the scRNA-seq gene count matrix and normalized together using NormalizeData function in Seurat.

### Cell-to-cell communication of scRNA-seq data

The CellPhoneDB algorithm was used to infer cell-to-cell interactions^63^. Briefly, the algorithm allows to detect ligand-receptor interactions between cell types in scRNA-seq data. We assessed the amount of interactions that are shared and specific for *i)* COVID-19 *versu*s non-COVID-19 and *ii)* mild *versus* critical COVID-19.

### Quantification and statistical analysis

Descriptive statistics are presented as median [interquartile range; IQR] (or median [range] if dataset contained only 2 variables) and n (%) for continuous and categorical variables, respectively. Statistical analyses were performed using R (version 3.6.3, R Foundation for Statistical Computing, R Core Team, Vienna, Austria). Statistical analyses were performed with a two-sided alternative hypothesis at the 5% significance level.

## Data availability

Raw sequencing reads of the scRNA-seq and scTCR-seq experiments generated for this study will be deposited in the EGA European Genome-Phenome Archive database. Based on SCope, which is an interactive web server for scRNA-seq data visualisation, a download of the read count matrix will be made available at http://blueprint.lambrechtslab.org. The publicly available datasets that supported this study are available from GEO GSE145926^7^, GEO GSE122960^10^ and from ArrayExpress E-MTAB-6149/E-MTAB-6653^9^.

## Additional resources

The findings outlined above are part of the COntAGIouS observational clinical trial: https://clinicaltrials.gov/ct2/show/NCT04327570.

## Acknowledgements

This project has received funding within the Grand Challenges Program of VIB. This VIB Program received support from the Flemish Government under the Management Agreement 2017-2021 (VR 2016 2312 Doc.1521/4). ADG acknowledges the financial support from Research Foundation Flanders (FWO) (G0B4620N; EOS grant: 30837538), KU Leuven (C14/19/098; POR/16/040) and Kom op Tegen Kanker (KOTK/2018/11509/1). L.V. is supported by an FWO PhD fellowship (grant number 11E9819N). P.V.M. is supported by an FWO PhD fellowship (grant number 1S66020N). E.W. is supported by Stichting tegen Kanker (Mandate for basic & clinical oncology research). J.W. is supported by an FWO Fundamental Clinical Mandate (1833317N).

We thank Dr. Wynand Van Rompaey, Dr. Julie Van Maercke, Dr. Nico De Crem, Dr. Sigurd Ghekiere and Dr. Thomas Demuynck for their help in patient recruitment and sample collection.

## Authors contributions

E.W., P.V.M., A.D.G., J.W., D.L. and J.Q. designed the experiments, developed the methodology, analysed and interpreted data and wrote the manuscript. S.J. and J.N. designed the experiments, developed the methodology and performed experiments. Y.V.H. and L.V. designed the experiments, collected and interpreted data. D.T., G.H., D.D., J.Y., J.G. and C.D. performed sample collection. A.B., B.B., B.M.D., P.M. and S.J. performed experiments and data analysis and interpretation. T.V.B., R.S., T.V.B. and E.H. provided technical assistance and performed experiments. E.W. and J.W. supervised the clinical study design and were responsible for coordination and strategy. All authors have approved the final manuscript for publication.

## CONTAGIOUS co-authors

Francesca Maria Bosisio, Michael Casaer, Frederik De Smet, Paul De Munter, Stephanie Humblet-Baron, Adrian Liston, Natalie Lorent, Kim Martinod, Paul Proost, Jeroen Raes, Karin Thevissen, Robin Vos, Birgit Weynand, Carine Wouters

## References

1. World Health Organization. WHO Coronavirus Disease (COVID-19) Dashboard. https://covid19.who.int/

2. Fu, L. et al. Clinical characteristics of coronavirus disease 2019 (COVID-19) in China: A systematic review and meta-analysis. J Infect. 80, 656–665 (2020).

3. Chen, G. et al. Clinical and immunological features of severe and moderate coronavirus disease 2019. J. Clin. Invest. 130, 2620–2629 (2020).

4. Liu, K. et al. Clinical characteristics of novel coronavirus cases in tertiary hospitals in Hubei Province. Chin. Med. J. (Engl). 133, 1025–1031 (2020).

5. Wen, W. et al. Immune cell profiling of COVID-19 patients in the recovery stage by single-cell sequencing. Cell Discov. 6, (2020).

6. Wilk, A. J. et al. A single-cell atlas of the peripheral immune response to severe COVID-19. Nat. Med. (2020).

7. Liao, M. et al. Single-cell landscape of bronchoalveolar immune cells in patients with COVID-19. Nat. Med. (2020).

8. Bost, P. et al. Host-viral infection maps reveal signatures of severe COVID-19 patients. Cell 1–14 (2020).

9. Lambrechts, D. et al. Phenotype molding of stromal cells in the lung tumor microenvironment. Nat. Med. 24, 1277–1289 (2018).

10. Reyfman, P. A. et al. Single-cell transcriptomic analysis of human lung provides insights into the pathobiology of pulmonary fibrosis. Am. J. Respir. Crit. Care Med. 199, 1517–1536 (2019).

11. Qian, J. et al. A Pan-cancer Blueprint of the Heterogeneous Tumour Microenvironment Revealed by Single-Cell Profiling. Cell Res. (2020).

12. Garg, A. D. & Agostinis, P. Cell death and immunity in cancer: From danger signals to mimicry of pathogen defense responses. Immunol. Rev. 280, 126–148 (2017).

13. Van Driel, B. J., Liao, G., Engel, P. & Terhorst, C. Responses to microbial challenges by SLAMF receptors. Front. Immunol. 7, 1–14 (2016).

14. Wherry, E. J. et al. Molecular Signature of CD8+ T Cell Exhaustion during Chronic Viral Infection. Immunity 27, 670–684 (2007).

15. Wang, W. H. et al. The role of galectins in virus infection - A systemic literature review. J. Microbiol. Immunol. Infect. (2019) doi:10.1016/j.jmii.2019.09.005.

16. Liu, W. et al. Tim-4 in Health and Disease: Friend or Foe? Front. Immunol. 11, 1–10 (2020).

17. Waite, J. C. & Skokos, D. TH17 response and inflammatory autoimmune diseases. Int. J. Inflam. 2012, (2012).

18. Chechlinska, M. et al. Molecular signature of cell cycle exit induced in human T lymphoblasts by IL-2 withdrawal. BMC Genomics 10, (2009).

19. Miyazaki, Y., Chen, L. C., Chu, B. W., Swigut, T. & Wandless, T. J. Distinct transcriptional responses elicited by unfolded nuclear or cytoplasmic protein in mammalian cells. Elife 4, 1–24 (2015).

20. Willingham, S. B. et al. The CD47-signal regulatory protein alpha (SIRPa) interaction is a therapeutic target for human solid tumors. Proc. Natl. Acad. Sci. U. S. A. 109, 6662–6667 (2012).

21. Garg, A. D., Romano, E., Rufo, N. & Agostinis, P. Immunogenic versus tolerogenic phagocytosis during anticancer therapy: Mechanisms and clinical translation. Cell Death Differ. 23, 938–951 (2016).

22. Cauwels, A., Rogge, E., Vandendriessche, B., Shiva, S. & Brouckaert, P. Extracellular ATP drives systemic inflammation, tissue damage and mortality. Cell Death Dis. 5, 1–7 (2014).

23. Krysko, D. V. et al. Emerging role of damage-associated molecular patterns derived from mitochondria in inflammation. Trends Immunol. 32, 157–164 (2011).

24. Riteau, N. et al. Extracellular ATP is a danger signal activating P2X7 receptor in lung inflammation and fibrosis. Am. J. Respir. Crit. Care Med. 182, 774–783 (2010).

25. Gavin, C. et al. The Complement System Is Essential for the Phagocytosis of Mesenchymal Stromal Cells by Monocytes. Front. Immunol. 10, (2019).

26. Tippett, E., Cameron, P. U., Marsh, M. & Crowe, S. M. Characterization of tetraspanins CD9, CD53, CD63, and CD81 in monocytes and macrophages in HIV-1 infection. J. Leukoc. Biol. 93, 913–920 (2013).

27. Lévesque, S. A., Kukulski, F., Enjyoji, K., Robson, S. C. & Sévigny, J. NTPDase1 governs P2X7-dependent functions in murine macrophages. Eur. J. Immunol. 40, 1473–1485 (2010).

28. Garg, A. D. et al. Pathogen response-like recruitment and activation of neutrophils by sterile immunogenic dying cells drives neutrophil-mediated residual cell killing. Cell Death Differ. 24, 832–843 (2017).

29. Galani, I. E. & Andreakos, E. Neutrophils in viral infections: Current concepts and caveats. J. Leukoc. Biol. 98, 557–564 (2015).

30. Laghlali, G., Lawlor, K. E. & Tate, M. D. Die another way: Interplay between influenza A virus, inflammation and cell death. Viruses 12, 1–23 (2020).

31. Zheng, S. et al. Viral load dynamics and disease severity in patients infected with SARS-CoV-2 in Zhejiang province, China, January-March 2020: Retrospective cohort study. BMJ 369, 1–8 (2020).

32. Dick, J. et al. C5a receptor 1 promotes autoimmunity, neutrophil dysfunction and injury in experimental anti-myeloperoxidase glomerulonephritis. Kidney Int. 93, 615–625 (2018).

33. Leitner, J., Herndler-Brandstetter, D., Zlabinger, G. J., Grubeck-Loebenstein, B. & Steinberger, P. CD58/CD2 Is the Primary Costimulatory Pathway in Human CD28 − CD8 + T Cells. J. Immunol. 195, 477–487 (2015).

34. Kaiser, B. K. et al. Interactions between NKG2x Immunoreceptors and HLA-E Ligands Display Overlapping Affinities and Thermodynamics. J. Immunol. 174, 2878–2884 (2005).

35. Schett, G. & Neurath, M. F. Resolution of chronic inflammatory disease: universal and tissue-specific concepts. Nat. Commun. 9, (2018).

36. Arandjelovic, S. & Ravichandran, K. S. Phagocytosis of apoptotic cells in homeostasis. Nat. Immunol. 16, 907–917 (2015).

37. Tay, M. Z., Poh, C. M., Rénia, L., MacAry, P. A. & Ng, L. F. P. The trinity of COVID-19: immunity, inflammation and intervention. Nat. Rev. Immunol. 20, 363–374 (2020).

38. Bratton, D. L. & Henson, P. M. Neutrophil clearance: When the party is over, clean-up begins. Trends Immunol. 32, 350–357 (2011).

39. Hochreiter-hufford, A. & Ravichandran, K. S. Clearing the Dead : Apoptotic Cell Sensing. Cold Spring Harb. Perspect. Biol. 5, a008748 (2013).

40. Merad, M. & Martin, J. C. Pathological inflammation in patients with COVID-19: a key role for monocytes and macrophages. Nat. Rev. Immunol. 20, 355–362 (2020).

41. Jose, R. J. & Manuel, A. COVID-19 cytokine storm: the interplay between inflammation and coagulation. Lancet Respir. Med. 8, e46–e47 (2020).

42. McGonagle, D., O’Donnell, J. S., Sharif, K., Emery, P. & Bridgewood, C. Immune mechanisms of pulmonary intravascular coagulopathy in COVID-19 pneumonia. Lancet Rheumatol. 2019, 1–9 (2020).

43. Wan, T., Zhao, Y., Fan, F., Hu, R. & Jin, X. Dexamethasone inhibits S. aureus-induced neutrophil extracellular pathogen-killing mechanism, possibly through toll-like receptor regulation. Front. Immunol. 8, (2017).

44. Cain, D. W. & Cidlowski, J. A. Immune regulation by glucocorticoids. Nat. Rev. Immunol. 17, 233–247 (2017).

45. Gutsol, A. A., Sokhonevich, N. A., Seledtsov, V. I. & Litvinova, L. S. Dexamethasone effects on activation and proliferation of immune memory T cells. Bull. Exp. Biol. Med. 155, 474–476 (2013).

46. Barnes, B. J. et al. Targeting potential drivers of COVID-19: Neutrophil extracellular traps. J. Exp. Med. 217, 1–7 (2020).

47. Uli, O. et al. Azithromycin modulates neutrophil function and circulating inflammatory mediators in healthy human subjects. Eur. J. Pharmacol. 450, 277–289 (2002).

48. Geleris, J. et al. Observational Study of Hydroxychloroquine in Hospitalized Patients with Covid-19. N. Engl. J. Med. 1–8 (2020).

49. Schrezenmeier, E. & Dörner, T. Mechanisms of action of hydroxychloroquine and chloroquine: implications for rheumatology. Nat. Rev. Rheumatol. 16, 155–166 (2020).

50. Grein, J. et al. Compassionate Use of Remdesivir for Patients with Severe Covid-19. N. Engl. J. Med. 2327–2336 (2020).

51. Stuart, T. et al. Comprehensive Integration of Single-Cell Data. Cell 177, 1888–1902.e21 (2019).

52. Van Den Brink, S. C. et al. Single-cell sequencing reveals dissociation-induced gene expression in tissue subpopulations. Nat. Methods 14, 935–936 (2017).

53. Buffa, F. M., Harris, A. L., West, C. M. & Miller, C. J. Large meta-analysis of multiple cancers reveals a common, compact and highly prognostic hypoxia metagene. Br. J. Cancer 102, 428–435 (2010).

54. McGinnis, C. S., Murrow, L. M. & Gartner, Z. J. DoubletFinder: Doublet Detection in Single-Cell RNA Sequencing Data Using Artificial Nearest Neighbors. Cell Syst. 8, 329–337.e4 (2019).

55. Wolock, S. L., Lopez, R. & Klein, A. M. Scrublet: Computational Identification of Cell Doublets in Single-Cell Transcriptomic Data. Cell Syst. 8, 281–291.e9 (2019).

56. Street, K. et al. Slingshot: Cell lineage and pseudotime inference for single-cell transcriptomics. BMC Genomics 19, (2018).

57. Zhu, W. et al. A high density of tertiary lymphoid structure B cells in lung tumors is associated with increased CD4+ T cell receptor repertoire clonality. Oncoimmunology 4, (2015).

58. Robinson, J. P. W. et al. The limitations of diversity metrics in directing global marine conservation. Mar. Policy 48, 123–125 (2014).

59. Aibar, S. et al. SCENIC: Single-cell regulatory network inference and clustering. Nat. Methods 14, 1083–1086 (2017).

60. Van den Berge, K. et al. Trajectory-based differential expression analysis for single-cell sequencing data. Nat. Commun. 11, 1–13 (2020).

61. Federico, A. & Monti, S. HypeR: An R package for geneset enrichment workflows. Bioinformatics 36, 1307–1308 (2020).

62. Kim, D. et al. The Architecture of SARS-CoV-2 Transcriptome. Cell 181, 914–921.e10 (2020).

63. Efremova, M., Vento-Tormo, M., Teichmann, S. A. & Vento-Tormo, R. CellPhoneDB: inferring cell–cell communication from combined expression of multi-subunit ligand–receptor complexes. Nat. Protoc. 15, 1484–1506 (2020).

