## Supplementary Figures S1-S6 and Tables S1 for "Discriminating Mild from Critical COVID-19 by Innate and Adaptive Immune Single-cell Profiling of Bronchoalveolar Lavages"

### Items Included

***Supplementary Figure Legends***

Fig. S1, related to Fig. 1

Fig. S2, related to Fig. 2

Fig. S3, related to Fig. 5

Fig. S4, related to Fig. 5

Fig. S5, related to Fig. 6 and Fig. 7

Fig. S6, related to Fig. 8

***Supplementary Table Legends***

Table S1, related to Fig. 1

Fig. S1

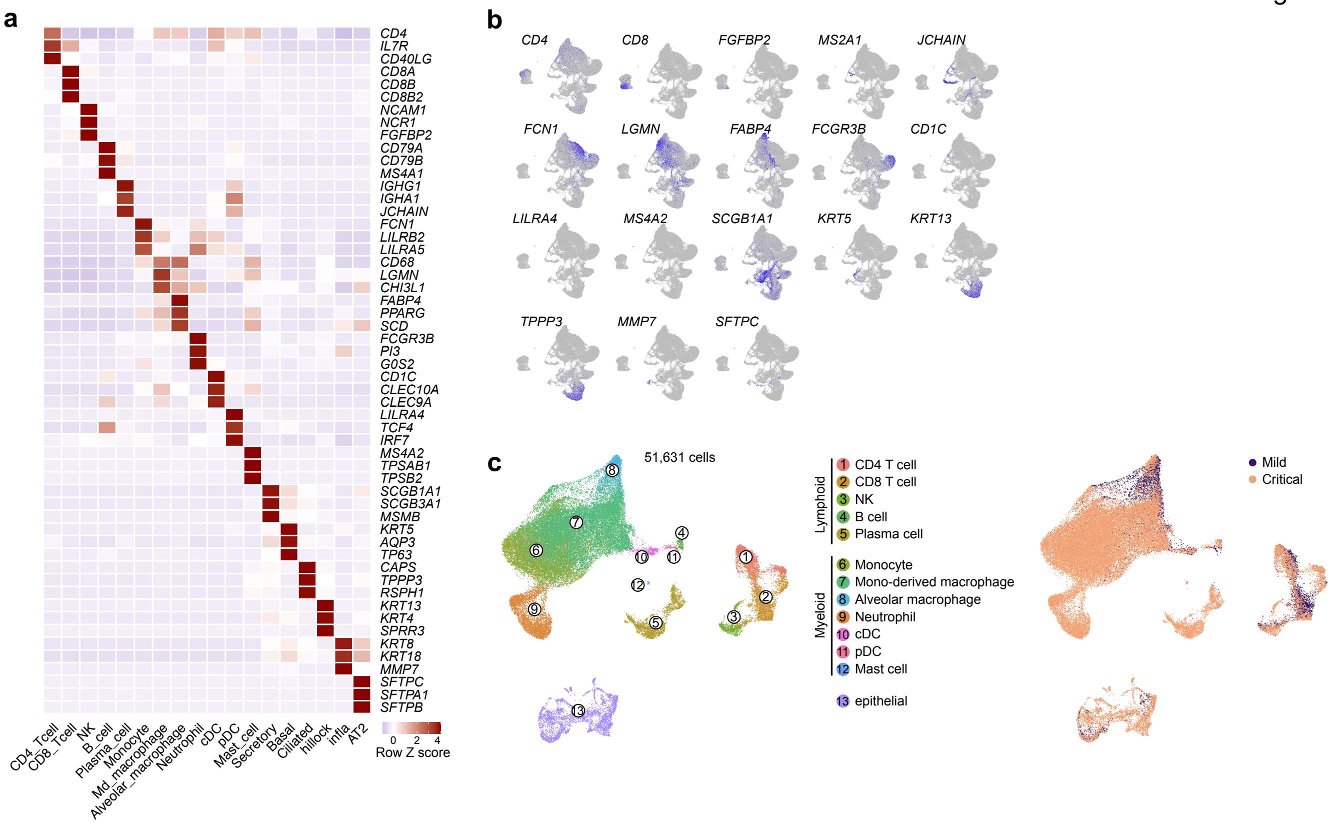
a
CD4
IL7R
CD40LG
CD8A
CD8B
CD8B2
NCAM1
NCR1
FGFBP2
CD79A
CD79B
MS4A1
IGHG1
IGHA1
JCHAIN
FCN1
LILRB2
LILRA5
CD68
LGMN
CHI3L1
FABP4
PPARG
SCD
FCGR3B
PI3
G0S2
CD1C
CLEC10A
CLEC9A
LILRA4
TCF4
IRF7
MS4A2
TPSAB1
TPSB2
SCGB1A1
SCGB3A1
MSMB
KRT5
AQP3
TP63
CAPS
TPPP3
RSPH1
KRT13
KRT4
SPRR3
KRT8
KRT18
MMP7
SFTPC
SFTPA1
SFTPB
CD4_Tcell
CD8_Tcell
NK
B_cell
Plasma_cell
Monocyte
Md_macrophage
Alveolar_macrophage
Neutrophil
cDC
pDC
Mast_cell
Secretory
Basal
Ciliated
hillock
infla
AT2
0 2 4
Row Z score
b
CD4
CD8
FGFBP2
MS2A1
JCHAIN
FCN1
LGMN
FABP4
FCGR3B
CD1C
LILRA4
MS4A2
SCGB1A1
KRT5
KRT13
TPPP3
MMP7
SFTPC
c
51,631 cells
Lymphoid
1 CD4 T cell
2 CD8 T cell
3 NK
4 B cell
5 Plasma cell
Myeloid
6 Monocyte
7 Mono-derived macrophage
8 Alveolar macrophage
9 Neutrophil
10 cDC
11 pDC
12 Mast cell
13 epithelial
Mild
Critical

**Fig. S1 Cell type annotation and clustering of scRNA-seq data by Liao et al.**

**a-b** Heatmap showing the expression of 3 marker genes (**a**) and feature plots showing 1 marker gene (**b**) for each of the different cell types. **c** UMAP showing 51,631 cells identified in scRNA-seq data by Liao et al., color-coded per cell type (left) and per disease severity status (right).

Fig. S2

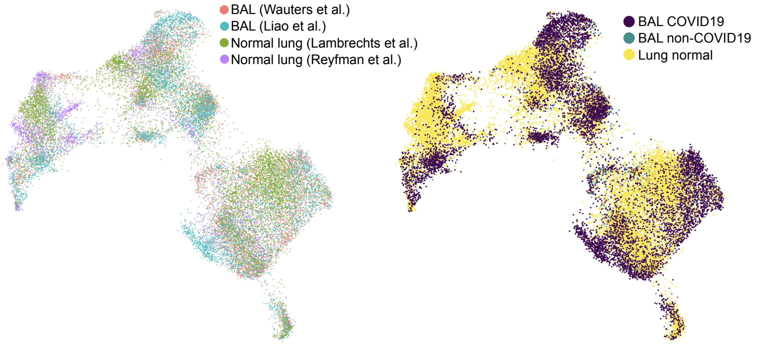

BAL (Wauters et al.)
BAL (Liao et al.)
Normal lung (Lambrechts et al.)
Normal lung (Reyfman et al.)
BAL COVID19
BAL non-COVID19
Lung normal

**Fig. S2 Single-cell profiling of T-/NK-cell phenotypes**

UMAP showing 23,468 T-/NK-cells color-coded per data origin (left) and per COVID-19 disease status (right).

Fig. S3

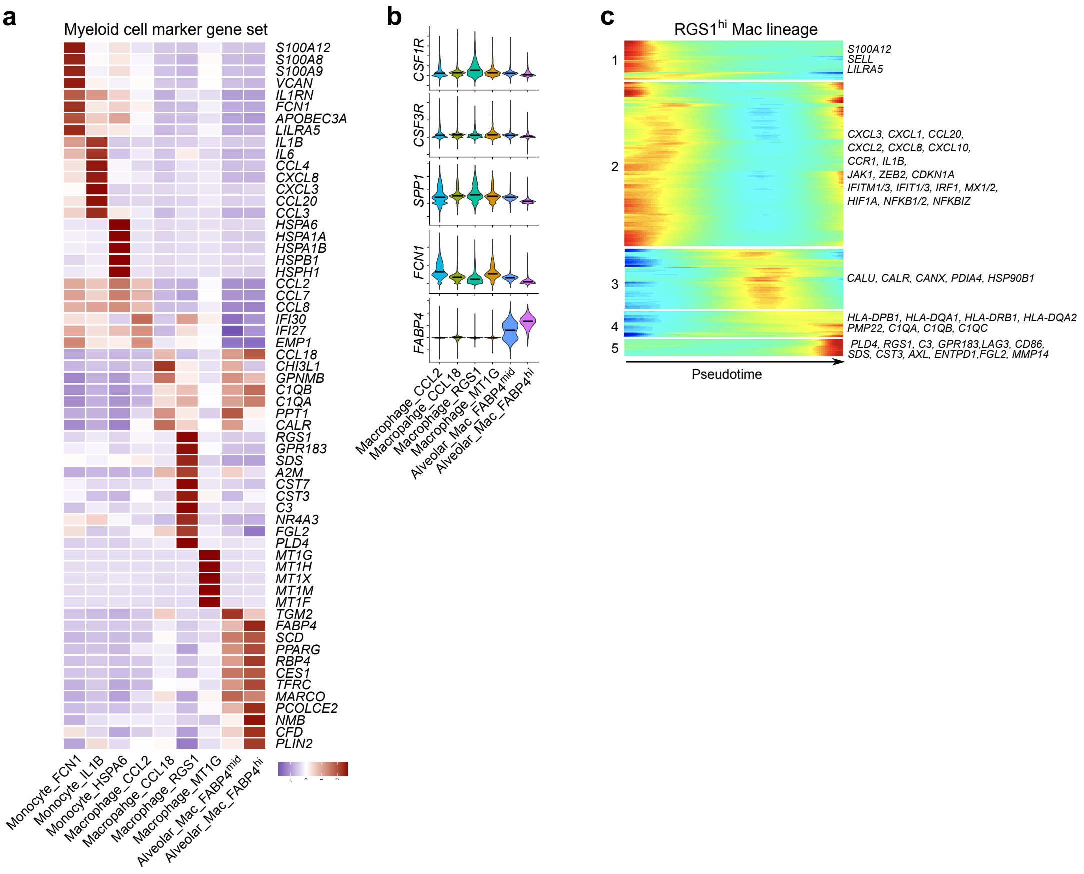
a
Myeloid cell marker gene set
S100A12
S100A8
S100A9
VCAN
IL1RN
FCN1
APOBEC3A
LILRA5
IL1B
IL6
CCL4
CXCL8
CXCL3
CCL20
CCL3
HSPA6
HSPA1A
HSPA1B
HSPB1
HSPH1
CCL2
CCL7
CCL8
IFI30
IFI27
EMP1
CCL18
CHI3L1
GPNMB
C1QB
C1QA
PPT1
CALR
RGS1
GPR183
SDS
A2M
CST7
CST3
C3
NR4A3
FGL2
PLD4
MT1G
MT1H
MT1X
MT1M
MT1F
TGM2
FABP4
SCD
PPARG
RBP4
CES1
TFRC
MARCO
PCOLCE2
NMB
CFD
PLIN2
Monocyte_FCN1
Monocyte_IL1B
Monocyte_HSPA6
Macrophage_CCL2
Macropahge_CCL18
Macrophage_RGS1
Macrophage_MT1G
Alveolar_Mac_FABP4mid
Alveolar_Mac_FABP4hi
b
CSF1R
CSF3R
SPP1
FCN1
FABP4
Macrophage_CCL2
Macropahge_CCL18
Macrophage_RGS1
Macrophage_MT1G
Alveolar_Mac_FABP4mid
Alveolar_Mac_FABP4hi
c
RGS1hi Mac lineage
1
S100A12
SELL
LILRA5
2
CXCL3, CXCL1, CCL20,
CXCL2, CXCL8, CXCL10,
CCR1, IL1B,
JAK1, ZEB2, CDKN1A
IFITM1/3, IFIT1/3, IRF1, MX1/2,
HIF1A, NFKB1/2, NFKBIZ
3
CALU, CALR, CANX, PDIA4, HSP90B1
4
HLA-DPB1, HLA-DQA1, HLA-DRB1, HLA-DQA2
PMP22, C1QA, C1QB, C1QC
5
PLD4, RGS1, C3, GPR183,LAG3, CD86,
SDS, CST3, AXL, ENTPD1,FGL2, MMP14
Pseudotime

**Fig. S3 Single-cell profiling of myeloid cells**

**a** Heatmap showing myeloid cell phenotypes with corresponding marker gene sets. **b** Violin plots of normalized marker gene expression, distinguishing 6 macrophage subclusters for their monocyte-derived characteristics, as indicated by monocyte lineage associated genes (*CSF1R*, *CSF3R*, *SPP1*, *FCN1*), and alveolar macrophage marker gene (*FABP4*). **c** Gene expression dynamics along the RGS1$^{high}$ macrophage lineage. Genes cluster into 5 gene sets, each of them characterized by specific expression profiles, as depicted by a selection of marker gene characteristic for each set.

Fig. S4

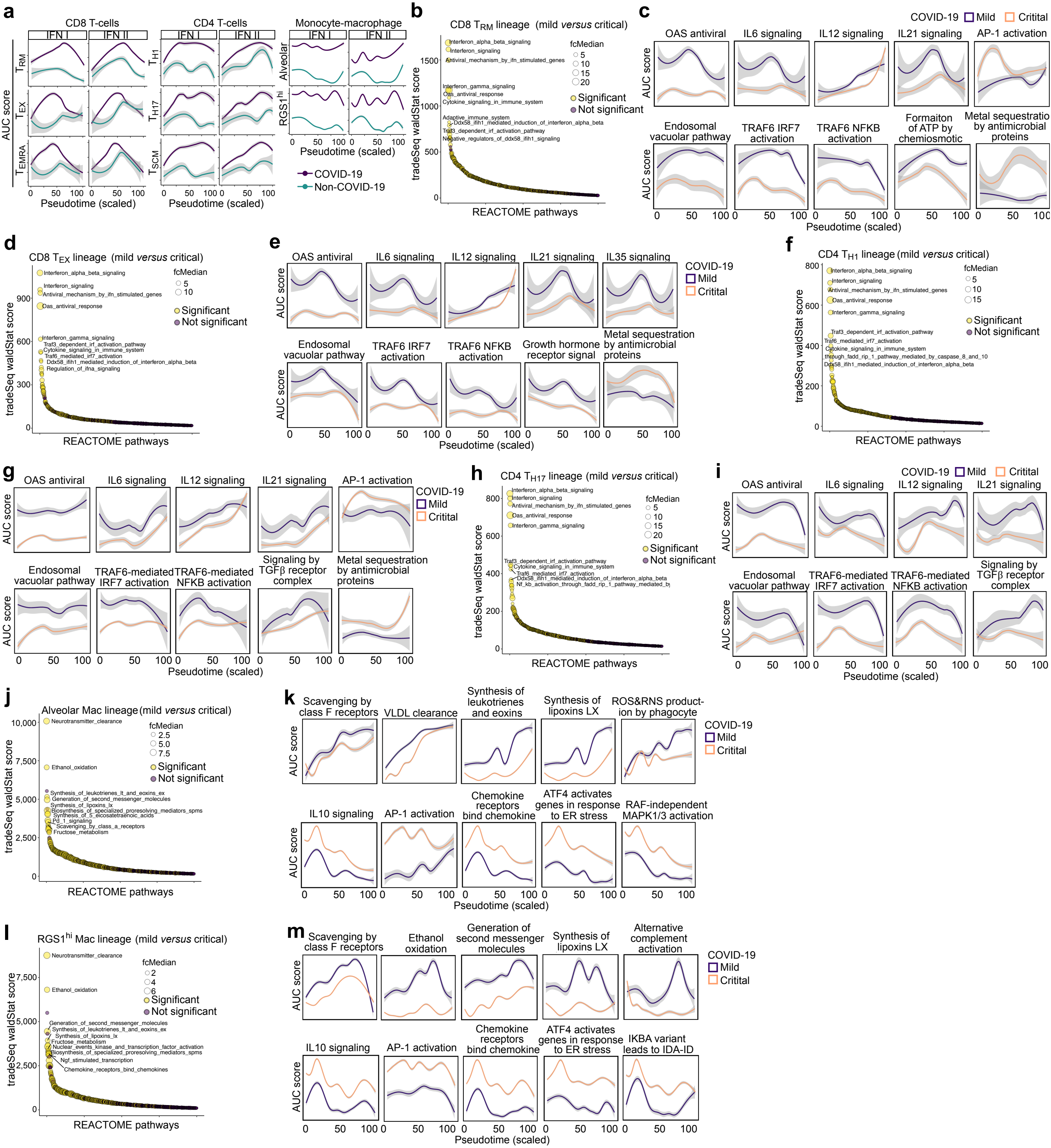
a
CD8 T-cells
CD4 T-cells
Monocyte-macrophage
IFN I
IFN II
AUC score
T_RM
T_EX
T_EMRA
T_H1
T_H17
T_SCM
Alveolar
RGS1^hi
Pseudotime (scaled)
COVID-19
Non-COVID-19
b
CD8 T_RM lineage (mild versus critical)
tradeSeq waldStat score
REACTOME pathways
fcMedian
Significant
Not significant
Interferon_alpha_beta_signaling
Interferon_signaling
Antiviral_mechanism_by_ifn_stimulated_genes
Interferon_gamma_signaling
Oas_antiviral_response
Cytokine_signaling_in_immune_system
Adaptive_immune_system
Ddx58_ifih1_mediated_induction_of_interferon_alpha_beta
Traf3_dependent_irf_activation_pathway
Negative_regulators_of_ddx58_ifih1_signaling
c
COVID-19
Mild
Critital
OAS antiviral
IL6 signaling
IL12 signaling
IL21 signaling
AP-1 activation
Endosomal vacuolar pathway
TRAF6 IRF7 activation
TRAF6 NFKB activation
Formaiton of ATP by chemiosmotic
Metal sequestration by antimicrobial proteins
AUC score
Pseudotime (scaled)
d
CD8 T_EX lineage (mild versus critical)
Interferon_alpha_beta_signaling
Interferon_signaling
Antiviral_mechanism_by_ifn_stimulated_genes
Oas_antiviral_response
Interferon_gamma_signaling
Traf3_dependent_irf_activation_pathway
Cytokine_signaling_in_immune_system
Traf6_mediated_irf7_activation
Ddx58_ifih1_mediated_induction_of_interferon_alpha_beta
Regulation_of_ifna_signaling
e
OAS antiviral
IL6 signaling
IL12 signaling
IL21 signaling
IL35 signaling
Endosomal vacuolar pathway
TRAF6 IRF7 activation
TRAF6 NFKB activation
Growth hormone receptor signal
Metal sequestration by antimicrobial proteins
f
CD4 T_H1 lineage (mild versus critical)
Interferon_alpha_beta_signaling
Interferon_signaling
Antiviral_mechanism_by_ifn_stimulated_genes
Oas_antiviral_response
Interferon_gamma_signaling
Traf3_dependent_irf_activation_pathway
Traf6_mediated_irf7_activation
Cytokine_signaling_in_immune_system
through_fadd_rip_1_pathway_mediated_by_caspase_8_and_10
Ddx58_ifih1_mediated_induction_of_interferon_alpha_beta
g
OAS antiviral
IL6 signaling
IL12 signaling
IL21 signaling
AP-1 activation
Endosomal vacuolar pathway
TRAF6-mediated IRF7 activation
TRAF6-mediated NFKB activation
Signaling by TGFβ receptor complex
Metal sequestration by antimicrobial proteins
h
CD4 T_H17 lineage (mild versus critical)
Interferon_alpha_beta_signaling
Interferon_signaling
Antiviral_mechanism_by_ifn_stimulated_genes
Oas_antiviral_response
Interferon_gamma_signaling
Traf3_dependent_irf_activation_pathway
Cytokine_signaling_in_immune_system
Traf6_mediated_irf7_activation
Ddx58_ifih1_mediated_induction_of_interferon_alpha_beta
i
OAS antiviral
IL6 signaling
IL12 signaling
IL21 signaling
Endosomal vacuolar pathway
TRAF6-mediated IRF7 activation
TRAF6-mediated NFKB activation
Signaling by TGFβ receptor complex
j
Alveolar Mac lineage (mild versus critical)
Neurotransmitter_clearance
Ethanol_oxidation
Synthesis_of_leukotrienes_lt_and_eoxins_ex
Generation_of_second_messenger_molecules
Synthesis_of_lipoxins_lx
Biosynthesis_of_specialized_proresolving_mediators_spms
Synthesis_of_5_eicosatetraenoic_acids
Pd_1_signaling
Scavenging_by_class_a_receptors
Fructose_metabolism
k
Scavenging by class F receptors
VLDL clearance
Synthesis of leukotrienes and eoxins
Synthesis of lipoxins LX
ROS&RNS production by phagocyte
IL10 signaling
AP-1 activation
Chemokine receptors bind chemokine
ATF4 activates genes in response to ER stress
RAF-independent MAPK1/3 activation
l
RGS1^hi Mac lineage (mild versus critical)
Neurotransmitter_clearance
Ethanol_oxidation
Generation_of_second_messenger_molecules
Synthesis_of_leukotrienes_lt_and_eoxins_ex
Synthesis_of_lipoxins_lx
Fructose_metabolism
Nuclear_events_kinase_and_transcription_factor_activation
Biosynthesis_of_specialized_proresolving_mediators_spms
Ngf_stimulated_transcription
Chemokine_receptors_bind_chemokines
m
Scavenging by class F receptors
Ethanol oxidation
Generation of second messenger molecules
Synthesis of lipoxins LX
Alternative complement activation
IL10 signaling
AP-1 activation
Chemokine receptors bind chemokine
ATF4 activates genes in response to ER stress
IKBA variant leads to IDA-ID

**Fig. S4 Pathway analysis along T-cell and monocyte-to-macrophage lineages**

**a** Differentially-activated IFN I and II pathways along the CD8$^+$, CD4$^+$ and monocyte-to-macrophage lineages, comparing COVID-19 *versus* non-COVID-19. **b** Differentially-activated REACTOME pathways for the CD8$^+$ $T_{RM}$-lineage, comparing mild *versus* critical COVID-19. **c** A selection of 10 differentially-activated REACTOME pathways along the CD8$^+$ $T_{RM}$-lineage, comparing mild *versus* critical COVID-19. **d** Differentially-activated REACTOME pathways along the CD8$^+$ $T_{EX}$-lineage, comparing mild *versus* critical COVID-19. **e** A selection of 10 differentially-activated REACTOME pathways along the CD8$^+$ $T_{ex}$-lineage, comparing mild *versus* critical COVID-19. **f** Differentially-activated REACTOME pathways for the CD4$^+$ $T_{H1}$-lineage, comparing mild *versus* critical COVID-19. **g** A selection of 10 differentially-activated REACTOME pathways along the CD4$^+$ $T_{H1}$-lineage, comparing mild *versus* critical COVID-19. **h** Differentially-activated REACTOME pathways for the CD4$^+$ $T_{H17}$-lineage, comparing mild *versus* critical COVID-19. **i** A selection of 8 differentially-activated REACTOME pathways along the CD4$^+$ $T_{H17}$-lineage, comparing mild *versus* critical COVID-19. **j** Differentially-activated REACTOME pathways for the alveolar macrophage lineage, comparing mild *versus* critical COVID-19. **k** A selection of 10 differentially-activated REACTOME pathways along the alveolar macrophage lineage, comparing mild *versus* critical COVID-19. **l** Differentially-activated REACTOME pathways for the RGS1-macrophage lineage, comparing mild *versus* critical COVID-19. **m** A selection of 10 differentially-activated REACTOME pathways along the RGS1-macrophage lineage, comparing mild *versus* critical COVID-19.

Fig. S5

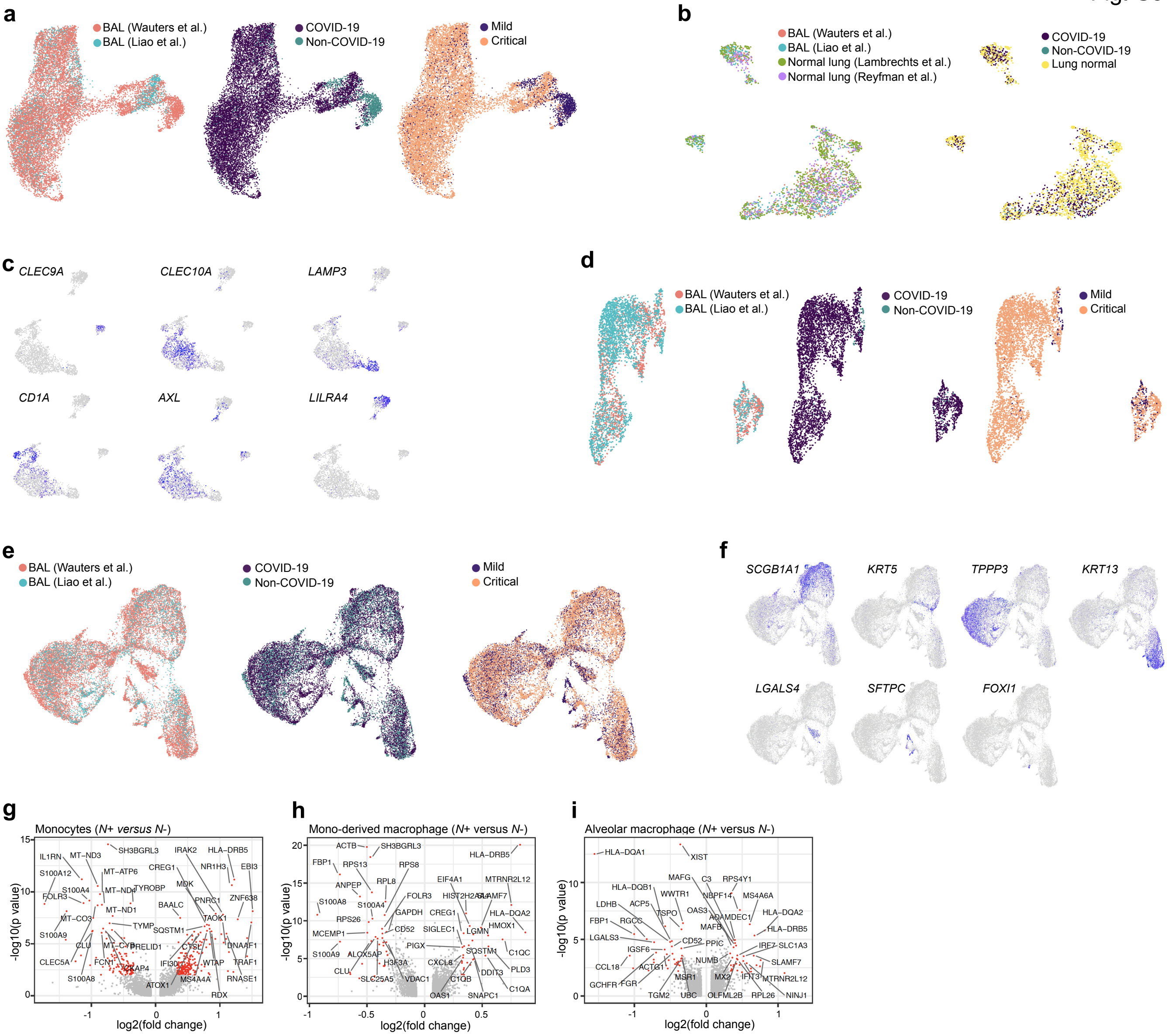
a
BAL (Wauters et al.)
BAL (Liao et al.)
COVID-19
Non-COVID-19
Mild
Critical
b
BAL (Wauters et al.)
BAL (Liao et al.)
Normal lung (Lambrechts et al.)
Normal lung (Reyfman et al.)
COVID-19
Non-COVID-19
Lung normal
c
CLEC9A
CLEC10A
LAMP3
CD1A
AXL
LILRA4
d
BAL (Wauters et al.)
BAL (Liao et al.)
COVID-19
Non-COVID-19
Mild
Critical
e
BAL (Wauters et al.)
BAL (Liao et al.)
COVID-19
Non-COVID-19
Mild
Critical
f
SCGB1A1
KRT5
TPPP3
KRT13
LGALS4
SFTPC
FOXI1
g
Monocytes (N+ versus N-)
-log10(p value)
log2(fold change)
h
Mono-derived macrophage (N+ versus N-)
-log10(p value)
log2(fold change)
i
Alveolar macrophage (N+ versus N-)
-log10(p value)
log2(fold change)

**Fig. S5 Neutrophil, dendritic, B- and epithelial cell phenotyping in COVID-19**

**a** UMAP showing 14,154 neutrophils color-coded per data origin (left), per disease status (middle) and per disease severity status (right). **b** UMAP of 1,410 DCs, color-coded per data origin (left) and per disease status (right). **c** Feature plots of marker genes, distinguishing 6 DC subclusters: type I classical dendritic cells (*CLEC9A*), type II classical dendritic cells (*CLEC10A*), migratory dendritic cells (*LAMP3*), Langerhans cell-like dendritic cells (*CD1A*), AS-DC (*AXL*) and plasmacytoid dendritic cells (*LILRA4*). **d** UMAP of 1,397 B-cells, color-coded per data origin (left), per disease status (middle) and per disease severity status (right). **e** UMAP of 22,215 epithelial cells, color-coded per data origin (left), per disease status (middle) and per disease severity status (right). **f** Feature plots of marker genes, distinguishing 7 epithelial cell subclusters: secretory cells (*SCGB1A1*), basal cells (*KRT5*), ciliated cells (*TPPP3*), hillock cells (*KRT13*), inflammatory cells (*LGALS4*), alveolar type II cells (*SFTPC*) and ionocytes (*FOXI1*). **g-i** Differentially expressed genes in *N*+ versus *N*- monocytes (**g**), monocyte-derived macrophages (**h**) and alveolar macrophages (**i**).

Fig. S6

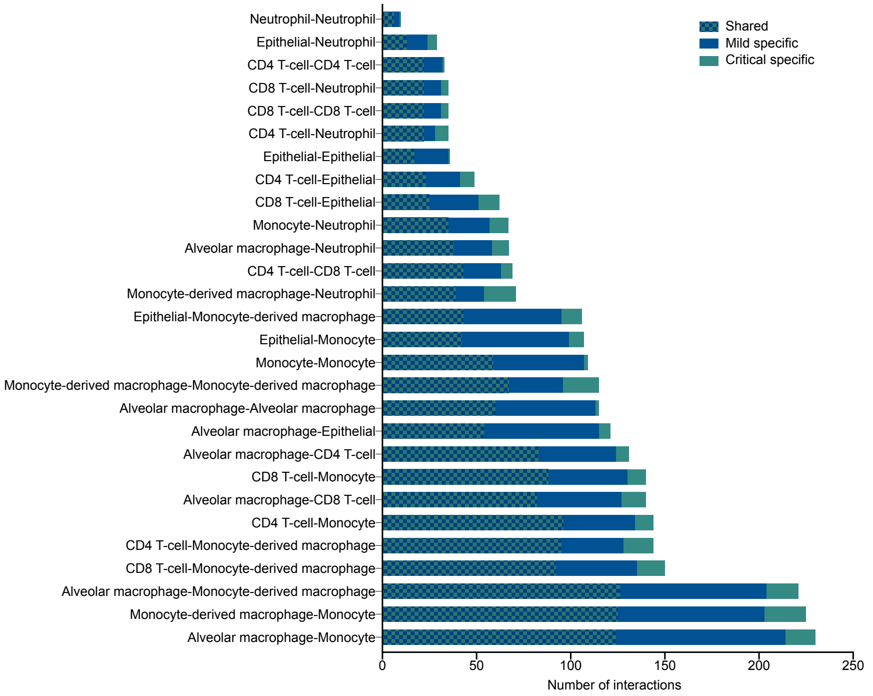

Neutrophil-Neutrophil
Epithelial-Neutrophil
CD4 T-cell-CD4 T-cell
CD8 T-cell-Neutrophil
CD8 T-cell-CD8 T-cell
CD4 T-cell-Neutrophil
Epithelial-Epithelial
CD4 T-cell-Epithelial
CD8 T-cell-Epithelial
Monocyte-Neutrophil
Alveolar macrophage-Neutrophil
CD4 T-cell-CD8 T-cell
Monocyte-derived macrophage-Neutrophil
Epithelial-Monocyte-derived macrophage
Epithelial-Monocyte
Monocyte-Monocyte
Monocyte-derived macrophage-Monocyte-derived macrophage
Alveolar macrophage-Alveolar macrophage
Alveolar macrophage-Epithelial
Alveolar macrophage-CD4 T-cell
CD8 T-cell-Monocyte
Alveolar macrophage-CD8 T-cell
CD4 T-cell-Monocyte
CD4 T-cell-Monocyte-derived macrophage
CD8 T-cell-Monocyte-derived macrophage
Alveolar macrophage-Monocyte-derived macrophage
Monocyte-derived macrophage-Monocyte
Alveolar macrophage-Monocyte
0
50
100
150
200
250
Number of interactions
Shared
Mild specific
Critical specific

**Fig. S6 Cell-to-cell interactions shared and specific in mild and critical COVID-19 BAL** Predicted number of cell-to-cell interactions between epithelial cell, neutrophil, monocyte, macrophage, $CD8^+$/$CD4^+$ T-cell from BAL of mild *versus* critical COVID-19.

### Supplementary Tables

#### Table S1. Demographics and characteristics of study cohort.

| | **Mild COVID-19 (n=2)** | **Critical COVID-19 (n=20)** | **Non-COVID pneumonia (n=13)** |
|---|---|---|---|
| Age, years | 65 [55-71] | 60 [54.5-68] | 68 [61-74] |
| Sex | .. | .. | .. |
| Men | 1 (50) | 15 (75) | 7 (54) |
| Women | 1 (50) | 5 (25) | 6 (46) |
| Medication history | .. | .. | .. |
| Immunosuppressants | 1 (50)[a] | 0 (0) | 4 (31)[b] |
| Time from illness onset to sampling (days) | 17.5 [15-20] | 19 [15.75-25] | 8 [2-14] |
| Bronchoalveolar lavage microbiology | .. | .. | .. |
| SARS-CoV-2 PCR positive | 2 (100) | 7 (35)[c] | 0 (0) |
| Other viral PCR positive | 0 (0) | 4 (20)[d] | 3 (23)[e] |
| Bacterial culture positive | 1 (50)[f] | 3 (15)[g] | 2 (15)[h] |
| PJP PCR positive | 0 (0) | 0 (0) | 5 (38) |
| Respiratory support | 2 (100) | 20 (100) | 8 (62) |
| Oxygen via nasal cannula | 2 (100) | 0 (0) | 7 (54) |
| Invasive ventilation | 0 (0) | 15 (75) | 1 (8) |
| Extracorporeal membrane oxygenation | 0 (0) | 5 (25) | 0 (0) |
| Medical treatment | 2 (100) | 20 (100) | 10 (77) |
| Antiviral therapy (<7d) | 0 (0) | 14 (70)[i] | 0 (0) |
| Antibiotics (<7d) | 2 (100) | 20 (100) | 8 (62) |
| Immunomodulatory therapy (<7d) | 0 (0) | 5 (25)[j] | 0 (0) |
| Fatal outcome[k] | 0 (0) | 1 (5) | 2 (15) |

**Legend to Supplementary Table S1:** Data are median [IQR], median [range] for the "Mild COVID-19" group or n (%). [a] 1 patient received chemotherapy. [b] 2 patients received chemotherapy, 1 patient taking abemaciclib and 1 patient taking mycophenolate mofetil/tacrolimus. [c] SARS-CoV-2 PCR not performed in 12 cases, 1 negative SARS-CoV-2 PCR on BAL (after initial PCR-confirmed diagnosis of COVID-19). [d] Herpes Simplex 1 PCR positive in 4 cases. [e] Herpes Simplex 1 PCR positive in 3 cases, 1 case of PCR-confirmed coronavirus HKU-1 co-infection. [f] 1 patient with positive *Escherichia coli* and *Pseudomonas aeruginosa* culture. [g] 3 patients with cultures positive for *Raoultella ornithinolytoca, Klebsiella aerogenes* or *Enterobacter cloacae*. [h] 2 patients with cultures positive for *Escherichia coli* or *Pseudomonas aeruginosa*; in 6 patients, organism remained unidentified, 4 of them received prior antibiotics. [i] Hydroxychloroquine administered in 12 cases, remdesivir administered in 2 cases. [j] 5 patients received >1mg/kg prednisone during 48h preceding sampling; 1 patient received Anakinra during the week preceding sampling. [k] Recruitment started on March 27th 2020, with outcome assessed on June 11th 2020. PJP: Pneumocystis jirovecii; SARS-CoV-2: Severe acute respiratory syndrome coronavirus 2.
